# Modulation of stress granule dynamics by phosphorylation and ubiquitination in plants

**DOI:** 10.1101/2024.02.20.581253

**Authors:** Siou-Luan He, Xiling Wang, Sungil Kim, Liang Kong, Lei Wang, Ping He, Libo Shan, Ying Wang, Jyan-Chyun Jang

**Author notes:** Correspondence: Jyan-Chyun Jang.

## Abstract

The Arabidopsis tandem CCCH zinc finger 1 (TZF1) is an RNA-binding protein that plays a crucial role in plant growth and stress response. TZF1 can localize to ribonucleoprotein (RNP) granules in response to various abiotic stresses. However, very little is known about the composition, function, and assembly mechanism of plant RNP granules. In this report, we show that TZF1 contains two intrinsically disordered regions (IDRs) necessary for its localization to stress granules (SGs), a subclass of RNP granules. TZF1 recruits mitogen-activated protein kinase (MAPK) signaling components and an E3 ubiquitin ligase KEEP-ON-GOING (KEG) to SGs. TZF1 is phosphorylated by MPKs and ubiquitinated by KEG. The phosphorylation sites of TZF1 were mapped by mass spectrometry. Mutant studies revealed that phosphorylation and ubiquitination of specific residues played differential roles in enhancing or reducing TZF1 SG assembly and protein-protein interaction with mitogen-activated kinase kinase 5 (MKK5) in SGs. TZF1 is extremely unstable, and its accumulation can be enhanced by proteosome inhibitor MG132. We showed that TZF1 was ubiquitinated in vivo and in vitro by KEG and TZF1 accumulated at a much lower level in gain-of-function mutant *keg-4*, compared to the WT. Ubiquitination appeared to play a positive role in TZF1 SG assembly, because either single or higher order mutations caused reduced number of SGs per cell, while enhanced the coalescence of small SGs into a large nucleus-like SG encompassing the nucleus. Together, our results demonstrate that the assembly of TZF1 SGs is distinctively regulated by ubiquitination and phosphorylation.

## Introduction

Ribonucleoprotein (RNP) granules are membrane-less biomolecular condensates normally formed through liquid-liquid phase separation (LLPS) driven by ATP and multivalent protein-protein, protein-RNA, and RNA-RNA interactions. The scaffold proteins in the RNP granules often contain intrinsically disordered domains (IDRs), low-complexity domains (LCDs), or prion-like domains (PrLDs) to facilitate the nucleation and growth of the condensates (1,2). Processing bodies (P-bodies, PBs) and stress granules (SGs) are two types of RNP granules found in eukaryotic cells. PBs and SGs play crucial roles in physiological and stress responses via dynamic regulation of signal transduction and mRNA metabolism. PBs and SGs provide a unique spatiotemporal regulatory mechanism that mediates various cellular processes (1,3–7). Earlier research suggests that PBs and SGs carry out distinct functions, given unique protein and RNA compositions are found in PBs and SGs, respectively. However, the boundaries between PBs and SGs have become blurred with more recent research (1). Many proteins have been found in both compartments, such as Argonaute 1/2 (Ago1/2), early Initiation Factor 4E (eIF4E), Apolipoprotein B mRNA-editing Enzyme Catalytic polypeptide 1-like 3G (APOBEC3G), and Tristertraprolin (TTP) in non-plant systems (8,9), and heat shock proteins and RNA helicases in plant system (2), suggesting overlapping functions and constant dynamic assembly and in some occasions protein exchange between PBs and SGs (1,10).

PBs are constitutive cytoplasmic RNP granules that consist of non-translating mRNAs, mRNA decay factors, translational repressors, and various RNA-binding proteins (RBPs) involved in mRNA storage, degradation, and translational repression (11). As in mammals, plant PBs contain conserved RNA degradation machineries, such as mRNA decapping factors (DCP1 and DCP2) and 5’-3’ processing exonucleases (XRN4) (9,12). Mutations in genes encoding essential components of PBs in Arabidopsis, such as DCP1 and DCP2, cause growth defects, suggesting an essential role of mRNA decapping in plant development (12). In addition, PB components are involved in both biotic and abiotic stress responses. For example, microbe-associated molecular patterns (MAMPs) were shown to modulate the dynamic interaction between DCP1, DCP2, and XRN4, assembly of PBs, and selective mRNA decay in plant immunity mediated by mitogen-activate protein kinase (MAPK) signaling cascade (13). The *Arabidopsis* DCP1 is phosphorylated by mitogen-activated protein kinase 6 (MPK6) and this process is critical for plant dehydration stress tolerance (14). However, whether MAPK components are recruited to PBs to phosphorylate DCP1 and if PBs are required for dehydration response remain to be addressed.

SGs are another class of cytoplasmic RNP granules that are transiently formed in response to various cellular stressors, such as heat shock, oxidative stress, viral infection, or nutrient deprivation (1,10,15). When cells encounter stress, translation initiation is often inhibited, leading to the accumulation of untranslated mRNAs. These untranslated mRNAs, along with various RBPs, are the main components for SG assembly. SGs help preserve mRNAs during stress and facilitate their translation after stress relief (Riggs et al., 2020). In mammals, SGs are typically formed by the aggregation of untranslated mRNAs, stalled translation initiation complexes, small ribosome subunits, and RBPs like T-cell-restricted intracellular antigen-1 (TIA-1) and Ras-GAP SH3 domain-binding proteins (G3BP1 and G3BP2), as well as many other proteins involved in signal transduction (16). In plants, the functions of SGs are less well characterized than in mammals, and the dynamics of compositional and functional changes of SGs in response to various cues is also under-investigated. Nevertheless, several plant SG proteins have been identified and characterized based on their homology with animal and yeast proteins or the results of proteomic studies (2). For instance, Tudor Staphylococcal Nuclease (TSN) proteins have been identified as a core component of plant SGs (17). The RNA-binding protein 47b (Rbp47b) (18) and oligouridylate binding protein 1B (UBP1B) (19) are the RBPs most closely related to mammalian TIA-1.

Plant tandem CCCH zinc finger proteins (TZFs) have been found in both PBs and SGs (20). TZFs are evolutionarily conserved in eukaryotes and they are characterized by a TZF motif consisting of two identical CCCH domains (C-X7-8-C-X5-C-X3-H) separated by 18 amino acids (21). However, a unique group of plant TZF proteins contain an arginine-rich (RR) region preceding a variant TZF motif consisting of two distinct CCCH domains (C-X_7-8_-C-X_5_-C-X_3_-H-X_16_ and C-X_5_-C-X_4_-C-X_3_-H) called RR-TZF proteins. Genes encoding RR-TZF proteins have been found in numerous higher plants, including *Arabidopsis* (TZF1-11) (22–26). Plant RR-TZF proteins participate in a plethora of biological processes including hormone-mediated growth and stress responses such as leaf senescence (OsTZF1 and OsTZF2) (27,28), ABA/GA-mediated growth and abiotic stress responses (TZF1) (29), seed germination (TZF4/5/6) (30), and flowering time (MsZFN) (31). The mammalian TZF protein tristetraprolin (TTP) is found in PBs and SGs and is participated in the posttranscriptional regulation of gene expression by binding to mRNAs (32). A classic model of TTP in mRNA regulation has been well established—TTP can trigger the decay of *Tumor Necrosis Factor-α* (*TNF-α*) mRNA by binding to its AU-rich elements (AREs) at 3’-UTR and recruiting deadenylation and decapping complexes to the substrate (33,34). In plants, TZF1/4/5/6/9 (23), OsTZF1/7 (28,35), and OsC3H10 (36) have been reported to colocalize with PBs and SGs markers. TZF1 can directly bind to U rich region of *Target of Rapamycin* (*TOR*) mRNA at 3’-UTR and trigger *TOR* mRNA degradation (37). OsTZF1 (28) and OsTZF7 (35) can bind ARE-like motifs within 3’-UTRs of downregulated target genes and likely induce mRNA turnover.

Although the biophysical mechanisms underpinning the assembly of biomolecular condensates via LLPS have been thoroughly investigated (1,5), the signal transduction mechanisms that trigger these processes are far from complete understood. Post-translational modifications (PTMs) play a crucial role in the regulation of SG assembly and disassembly. The dynamic nature of SG assembly is closely tied to the PTM status of the scaffold protein components. Alterations in PTMs can impact the formation, stability, and dissolution of SGs in response to cellular stress (38–40). Phosphorylation is a common PTM that regulates SG dynamics via the impacts on SG protein components. For example, phosphorylation of TTP by MAPKAP kinase-2 (MK2) promotes its binding to 14-3-3 adaptor proteins, thereby excluding TTP from SGs and stabilizing the ARE-containing target mRNAs (41). In plants, bacterial flagellin or flg22 peptides induces *Arabidopsis* TZF9 phosphorylation via two MAMP-responsive MPKs, MPK3 and MPK6. Phosphorylation of TZF9 diminishes cytoplasmic granules and RNA-binding properties (42). In addition, ubiquitination is one of the PTMs that marks proteins for degradation or regulates protein activity. Ubiquitin ligases and deubiquitinating enzymes can influence SGs dynamics by modifying the ubiquitination status of key SG proteins (43,44). The ubiquitination of some stress granule components may target them for degradation, leading to SGs disassembly. For examples, G3BP1 undergoes K63-linked ubiquitylation in the disassembly of SGs formed under heat stress (45). Two SG proteins carrying ubiquitin associated (UBA) domains, UBAP2L and UBQLN2, have been found to regulate SG assembly, but their roles are not dependent on the UBA domain (46,47). These results suggest a role for ubiquitination in regulating SG disassembly, but its impact on SG assembly remains unclear.

In this study, we have demonstrated that Arabidopsis TZF1 is a SG resident protein. Deletion of either or both IDRs flanked the RR-TZF motif could almost eliminate TZF1 SG assembly completely. TZF1 could interact with MAPK signaling cascade components in SGs. TZF1 recruits MPKs to SGs and is phosphorylated by MPK3/6. The potential phosphorylation sites of TZF1 were mapped by mass spectrometry in the absence/presence of a potent MAMP—flg22. Analysis of site-directed single and higher order mutations of potential phosphorylation sites revealed that phosphorylation of specific residues played differential roles in enhancing or reducing SG assembly and protein-protein interaction with an MPK3/6 upstream kinase—mitogen-activated kinase kinase 5 (MKK5) in SGs. Mutant analysis also identified two potential 14-3-3 adaptor protein binding sites to be critical for TZF1 SG assembly and protein-protein interaction with MKK5 in SGs. For the role of ubiquitination, TZF1 protein accumulated at a lower level in a ubiquitin E3 ligase gain-of-function mutant *keg-4* and TZF1 was ubiquitinated by recombinant KEEP-ON-GOING (KEG) *in vitro*. Remarkably, ubiquitination played a positive role in SG assembly, because single or higher order mutations on predicted ubiquitination sites of TZF1 reduced the number of SGs per cell, while enhanced the coalescence of small SGs into a large single SG attaching to the nucleus. Together, our results demonstrate that the assembly of TZF1 into SGs is mediated by a wide array of post-translational modification mechanisms, in which ubiquitination and phosphorylation play a distinct role.

## Results

### TZF1 is an SG component

We have shown previously that TZF1 could co-localize with both PB (DCP2) and SG (PABP8) markers (48). However, DCP2 is not a PB-specific marker (49). We therefore re-examined the sub-cellular localization of TZF1 using a set of different markers. The TZF1-GFP fusion protein is functional as reported recently (50). Results showed that TZF1 could only very partially co-localize with PB marker DCP1, but completely co-localize with SG marker UBP1b (51) (Figure 1A). IDRs are the key drives to trigger the assembly of biomolecular condensates within cells (1,2). These condensates play critical roles in cellular organization, signaling, and gene regulation, and abnormal condensate assemblies have been implicated in various diseases (5). The mammalian TZF1 homolog tristetraprolin (TTP) is both an RBP and an IDR scaffold protein for SG assembly (41). Using SMART algorithm (http://smart.embl-heidelberg.de/), two IDRs were identified in TZF1 protein. Deletion constructs of TZF1_ΔIDR1_ (aa 70-85, upstream of RR), TZF1^ΔIDR2^ (aa 218-233, downstream of TZF), and TZF1^ΔIDR1,2^ were then made accordingly (Figure 1B). Remarkably, deletion of either or both IDRs strongly reduced TZF1 SG assembly (Figures 1B-C), despite the truncated proteins were accumulated at the similar levels to that of the WT (Supplementary Figure 1A). As it was shown that RR-TZF domain of TZF1 protein is required for high-affinity RNA binding (52), the effects of TZF1^ΔRR^, TZF1^ΔTZF^, and TZF1^RR-TZF^ (Figure 1B) on TZF1 SG assembly were examined. Deletion of either RR or TZF caused severe reduction of TZF1 SG assembly (Figures 1B and D). The RR-TZF fragment alone conferred strong SG assembly (Figures 1B and D), suggesting that both or either N-or C-terminus contains negative elements for SG assembly. Results of immunoblot analysis indicated that reduced TZF1 SG granule assembly caused by ΔRR and ΔTZF was not due to reduced protein accumulation (Supplementary Figure 1B), indicating that TZF1 SG assembly is mediated through post-translational regulatory mechanisms.

**Figure 1.**
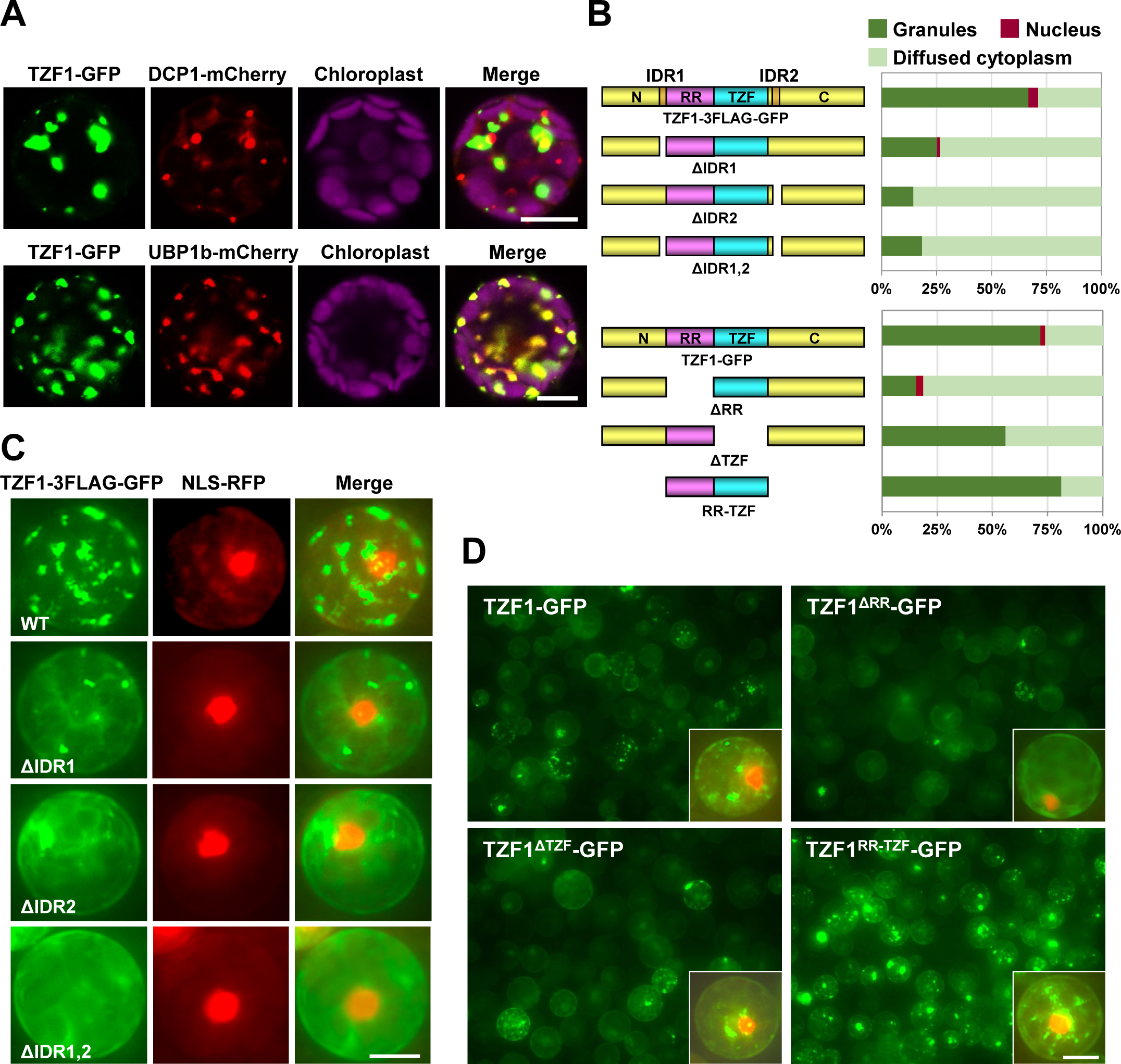
TZF1 is a stress granule component. **(A)** Confocal imaging showing that TZF1 is localized in cytoplasmic condensates and partially co-localized with PB marker DCP1, whereas completely co-localized with SG marker UBP1b. Scale bar= 10 μm. **(B)** The intrinsically disordered regions (IDR1 and IDR2) and RR-TZF motif are required for TZF1 cytoplasmic granule localization. Schematic representation of DNA constructs with deletion of predicted IDR (ΔIDR) and RR-TZF motif (ΔRR or ΔTZF) and corresponding quantitative analysis of TZF1 subcellular localization patterns. **(C)** TZF1 cytoplasmic granules were significantly reduced with IDR deletions in *Arabidopsis* protoplasts. Scale bar= 10 μm. **(D)** TZF1 cytoplasmic granules were reduced by the deletion of RR or TZF, but increased when only RR-TZF was present. Image in the insert is a single cell co-expressed with TZF1-GFP and NLS-RFP. Scale bar= 10 μm.

### TZF1 interacts with MAPK signaling components

To further explore the interacting proteins in the TZF1 protein complex, immunoprecipitation coupled with mass spectrometry (IP-MS) was performed using transgenic plants ectopically expressing *CaMV35S:TZF1-GFP* (29). Notably, two upstream kinases of mitogen-activated protein kinases (MKK4 and MKK5), two 14-3-3 adaptor proteins (At1g78300 and At5g38480), an E3 ubiquitin ligase KEG, and a conserved SG marker DEAD-box containing RNA helicase (RH6/8/12) were among the candidates identified by IP-MS (Figure 2A). Coincidentally, KEG was found to ubiquitinate MKK4 and MKK5 in modulating plant immunity (53). As we demonstrated previously that MAP kinase cascade (e.g., MPK3/6 and MKK4/5) is involved in MAMPs orchestrated PB dynamics (13), additional analyses were conducted to validate TZF1 protein-protein interaction with MAPK signaling components. Results showed that TZF1 could interact with MPK3, MPK6, MKK4, and MKK5 in both yeast-two-hybrid (Y-2-H) (Figure 2B) and co-immunoprecipitation (Co-IP) (Figure 2C) analyses. Consistent with TZF1’s sub-cellular localization, TZF1-MPK3, TZF1-MPK6, TZF1-MKK4, and TZF1-MKK5 protein complexes were very partially co-localized with PB marker DCP1, but completely co-localized with SG marker UBP1b in bi-molecular fluorescence complementation (BiFC) analyses (Supplementary Figure 2). Neither TZF1-nYFP nor TZF1-cYFP could interact with its corresponding BiFC empty vector construct. However, the MPK3-cYFP, MPK6-cYFP, and MKK4-cYFP did produce very weak nuclear signals with the nYFP empty vector. The MKK5-cYFP and nYFP empty vector generated unexpected visible YFP signals in the nuclei but never occurred in the cytoplasmic granules (Supplementary Figure 3A). To further confirm the specificity of the BiFC results, additional negative controls were included. Results showed that while MKK5-cYFP interacted with TZF1-nYFP in the cytoplasmic granules, it could not interact with five other nYFP fusion proteins (Supplementary Figure 3B), indicating the specificity of TZF1 interaction with MAPK signaling components (Supplementary Figure 2).

**Figure 2.**
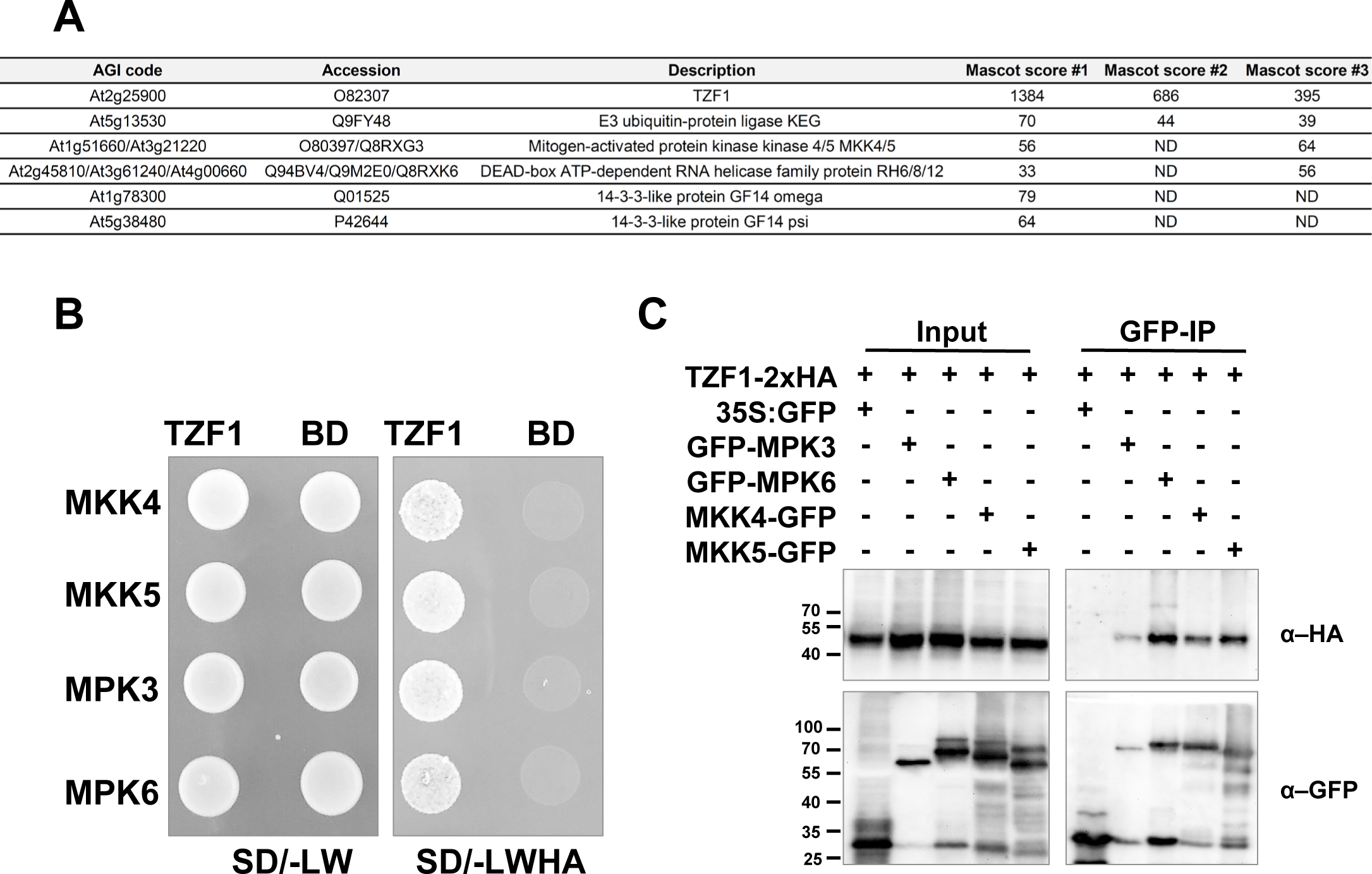
TZF1 interacts with mitogen activated kinase (MAPK) signaling cascade components. **(A)** Selected TZF1 protein complex components identified by immunoprecipitation coupled mass spectrometry. **(B)** TZF1 interacts with MAPK signaling components MKK4, MKK5, MPK3, and MPK6 in a Y-2-H assay, as indicated by the yeast growth on the quadruple amino acids dropout (-LWHA) selection plate. BD: empty vector with GAL4 binding domain was used as a negative control. **(C)** Co-IP analysis results indicate that TZF1 interacts with MPK3, MPK6, MKK4, and MKK5. Arabidopsis protoplasts were co-expressed with indicated constructs and IP was performed using anti-GFP antibody and Western blot was carried out using anti-HA and anti-GFP antibody, respectively.

The BiFC results prompted us to examine sub-cellular localization of MAPK signaling components. Using various fluorescence protein markers, we found that MKK4 and MKK5 were mainly localized in the cytoplasmic condensates, whereas MPK3 and MPK6 were mainly localized in the nucleus and cytoplasm (Figure 3A). Consistently, the MKK4-GFP and MKK5-GFP were localized in the cytoplasmic condensates in transgenic plants, albeit the nuclear signals were also quite visible (Figure 3B). Interestingly, individual MPK3, MPK6, MKK4, and MKK5 were not co-localized with PB marker DCP1, but completely co-localized with SG marker UBP1b (Supplementary Figure 4). Given TZF1-MPK3, TZF1-MPK6, TZF1-MKK4, and TZF1-MKK5 were colocalized with PB marker very partially and SG marker completely (Supplementary Figure 2), these results raise a possibility that MAPK cascade components are normally localized in nucleus, cytoplasm, or SGs to a lesser extent. MPK3/6 and MKK4/5 are recruited mainly to SGs when interacting with TZF1 (Figure 3C).

**Figure 3.**
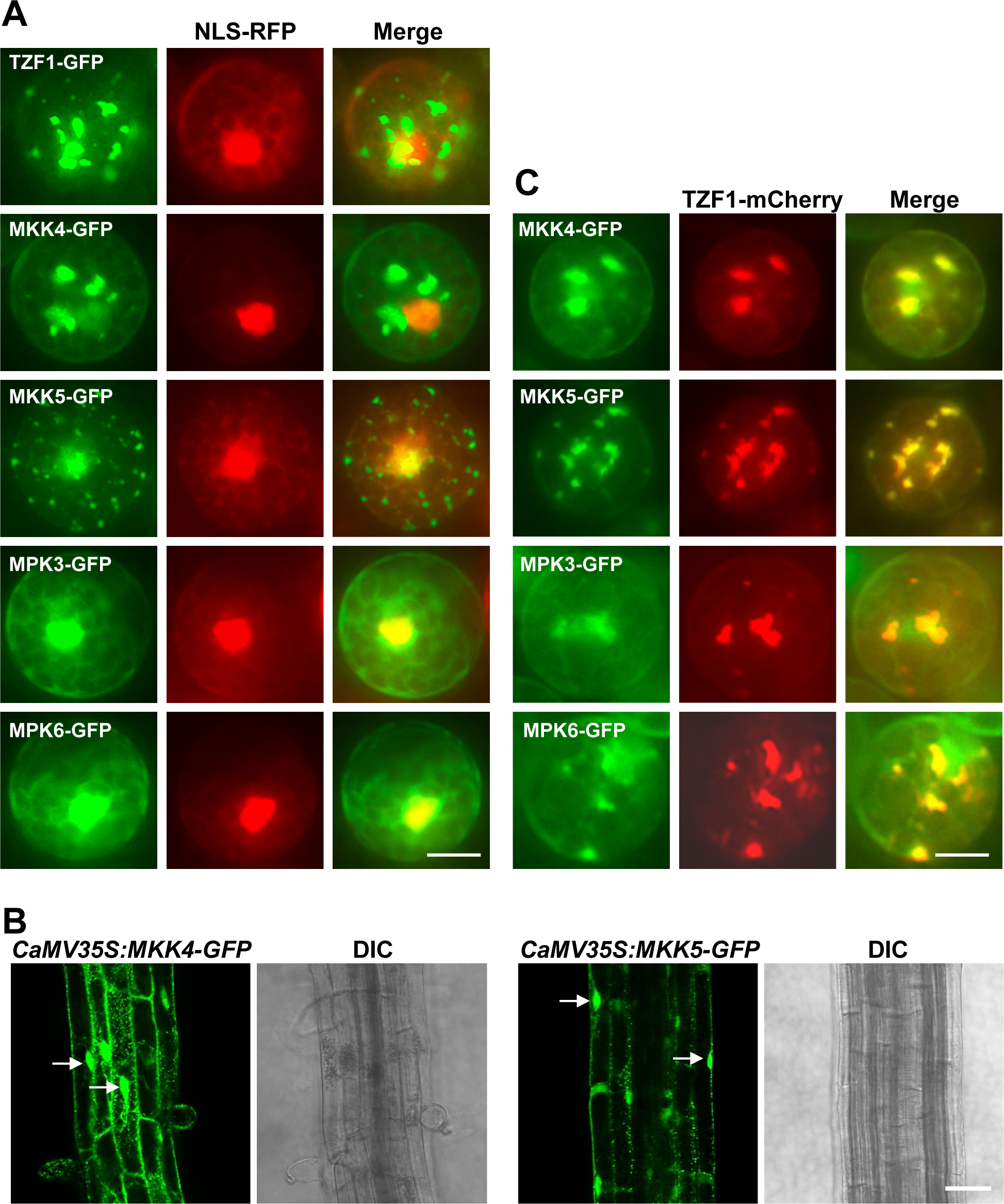
Subcellular localization of TZF1 and MAPK signaling components. **(A)** TZF1, MKK4, and MKK5 are localized in cytoplasmic condensates, whereas MPK3 and MPK6 are mainly localized in the nucleus in *Arabidopsis* protoplasts. NLS-RFP, a marker for nuclear proteins. Scale bar= 10 μm. **(B)** MKK4 and MKK5 are localized in cytoplasmic condensates in stable transgenic *Arabidopsis* plants. Shown are root tissues with GFP signals in cytoplasmic condensates throughout the cells and in the nuclei (arrows). Scale bar= 10 μm. **(C)** MPK3, MPK6, MKK4, and MKK5 co-localize with TZF1 in *Arabidopsis* protoplasts. Scale bar= 10 μm.

### TZF1 is phosphorylated by MPK3/6

The interaction between TZF1 and MAPK signaling components prompted us to verify if TZF1 can be phosphorylated by MPKs. Results of Phos-tag SDS-PAGE analysis indicated that TZF1 could be phosphorylated by MPK3, MPK4, and MPK6 upon flg22 treatment (Figure 4A). The human TZF homolog (TTP) is known to be heavily phosphorylated in numerous sites (54) and the phosphorylation status affects its subcellular localization, stability, and function (55). Furthermore, AtTZF9 is phosphorylated by MPK3 and MPK6 (56). We then performed phosphosite mapping by liquid chromatography coupled with tandem mass spectrometry (LC-MS/MS) to identify potential phosphorylation sites in TZF1. The eight identified phosphopeptides corresponded to ten residues S71, S73, S74, S80, S106, T110, S254, S266, T276, and S296 in TZF1 (Figures 4B-C, and 5). Among which, four sites (S71, S73, S80, and S254) were phosphorylated in the presence of flg22, with S80 showing the highest probability score and it was also predicted as a conserved MPK phosphosite with a signature SP motif (Figure 4C). The S254 was within a predicted 14-3-3 adaptor protein interacting site (results not shown) and adjacent to a predicted MPK phosphorylation site S255 (Figure 5). Three sites (S74, T110, and S296) appeared to be de-phosphorylated in the presence of flg22, with S74 showing the highest probability score. This might be of interesting because S74 clustered with three other flg22-induced phosphorylation sites S71, S73, and S80 (Figure 4C). The phosphorylation status of tree additional sites (S106, S266, and T276) did not seem to be affected by flg22 treatment. Notably, S106 and T276 were within a predicted 14-3-3 adaptor protein interacting site, respectively (Figure 4C).

**Figure 4.**
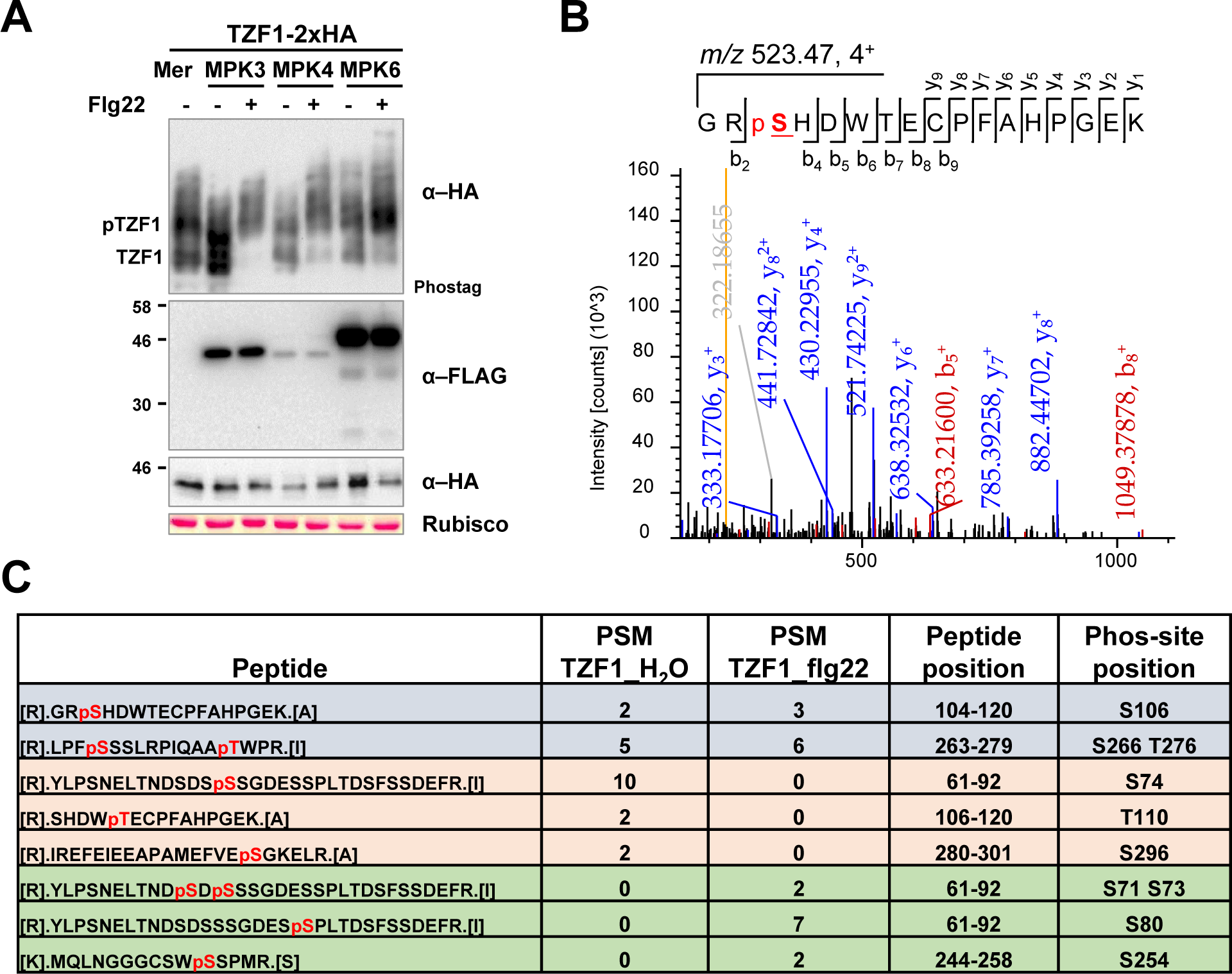
Flg22 induces phosphorylation of TZF1 on multiple serine and threonine residues. **(A)** Flg22-activated MPK3, MPK4, and MPK6 phosphorylate TZF1 in *Arabidopsis* protoplasts. Protoplasts co-expressing TZF1-HA with MPK3-FLAG, MPK4-FLAG, or MPK6-FLAG were treated with or without 0.1 μM flg22 for 15 min. Total proteins were separated with Mn^2+^-Phos-tag and regular SDS-PAGE gels, followed by immunoblot analysis with α-HA or α-FLAG antibodies. Protein loading is shown by Ponceau S staining for Rubisco. **(B)** LC-MS/MS spectrum of a phosphorylated peptide containing Ser-106 in TZF1 (TZF1^S106^). Protoplasts expressing TZF1-HA were treated without (H_2_O) or with 0.1 μM flg22 for 10 min. TZF1-HA was immunoprecipitated with α-HA magnetic beads and separated by SDS-PAGE gel, followed by digestion with trypsin and LC-MS/MS analysis to identify TZF1 phosphorylation sites. **(C)** List of TZF1 phosphorylation peptides identified by LC-MS/MS analysis. The peptide-spectrum match (PSM) indicates the number of identified phosphorylated peptides. TZF1^S74^, TZF1^T110^, and TZF1^S296^ were only identified in the H_2_O sample. TZF1^S71^, TZF1^S73^, TZF1^S80^, and TZF1^S254^ were only identified in the flg22-treated sample. TZF1^S106^, TZF1^S266^, and TZF1^T276^ were identified in both H_2_O and flg22-treated samples.

**Figure 5.**
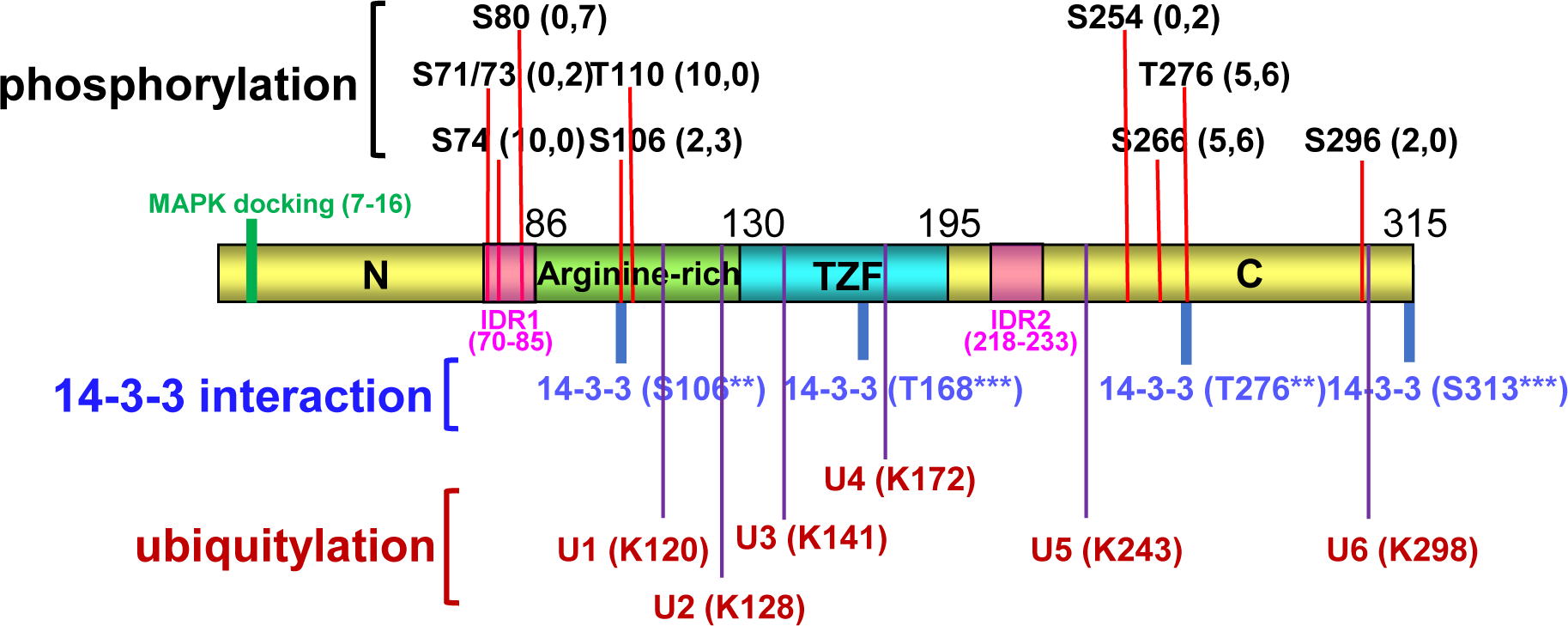
Schematic representation of domain structures and predicted post-translational modifications of TZF1. TZF1 is roughly divided into the N-terminus (N), arginine-rich motif (RR), tandem CCCH zinc finger motif (TZF), and C-terminus (C). The predicted MAPK docking site (aa 7 to 16) and two intrinsically disordered domains (IDR1 and IDR2) are also shown. Residues in black are predicted phosphorylation sites revealed by LC/MS-MS analysis. The two numbers in the parentheses next to the indicated residue are the peptide spectrum match (PSM) scores for sample treated without and with flg22, respectively. Residues in blue are potential 14-3-3 protein-protein interaction sites predicted by 14-3-3-Pred algorithm (https://www.compbio.dundee.ac.uk/1433pred). Residues in red are potential ubiquitylation sites predicted by BDM-PUB (Computational Prediction of Protein Ubiquitination Sites with a Bayesian Discriminant Method).

It has long been documented that phosphorylation of TTP by P38^MAPK^-MK2 signaling cascade prevents TTP localization to SGs and triggers protein-protein interaction between TTP and 14-3-3 protein (55,57). We therefore mutated S and T residues to A (phosphor-dead) or D (phosphor-mimetic) on the putative phosphorylation sites by site-directed mutagenesis to generate TZF1-3FLAG-GFP and TZF1-nYFP (BiFC) construct for sub-cellular localization and protein-protein interaction analysis, respectively. For the TZF1(WT)-3FLAG-GFP, the majority (∼70%) of the cells showed typical cytoplasmic granule (SG) pattern (Figure 6A #1). A small percentage of the cells displayed diffused cytoplasmic (Figure 6A #2), nuclear (Figure 6A #3) or nucleus-like (Figure 6A #4) pattern. It was intriguing that cells with reduced number of SGs appeared to be correlated with the formation of a large, coalesced SG closely associated with the nucleus (Figure 6A #4). Confocal microscopy rotating view of the large SG-nucleus complex revealed that the two organelles were physically attached (Supplementary Figure 5). Notably, the large coalesced TZF1 granules were completely co-localized with the SG marker UBP1b (Figure 6B), illustrating the dynamic assembly of TZF1 SGs in plant cells (1).

**Figure 6.**
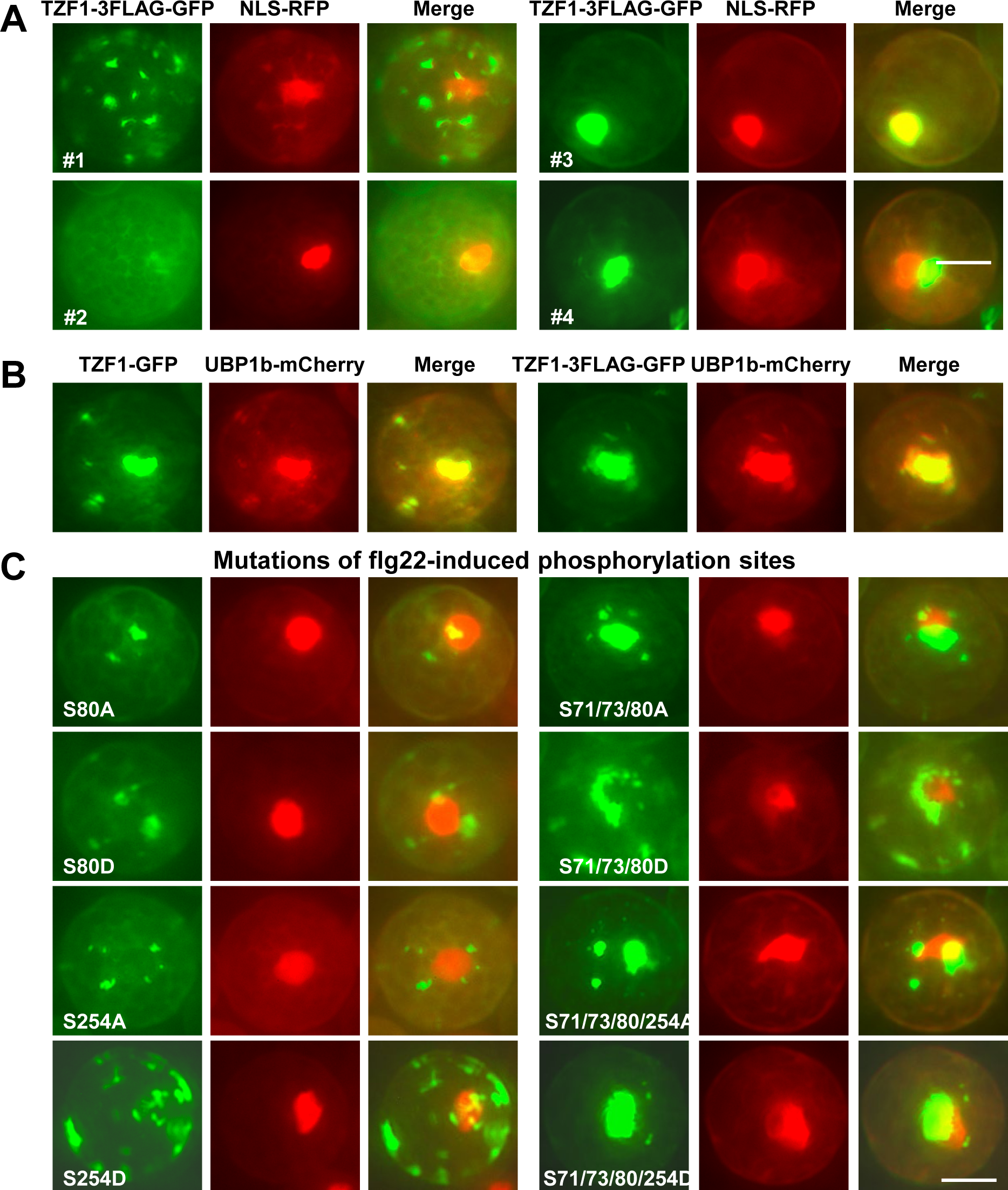
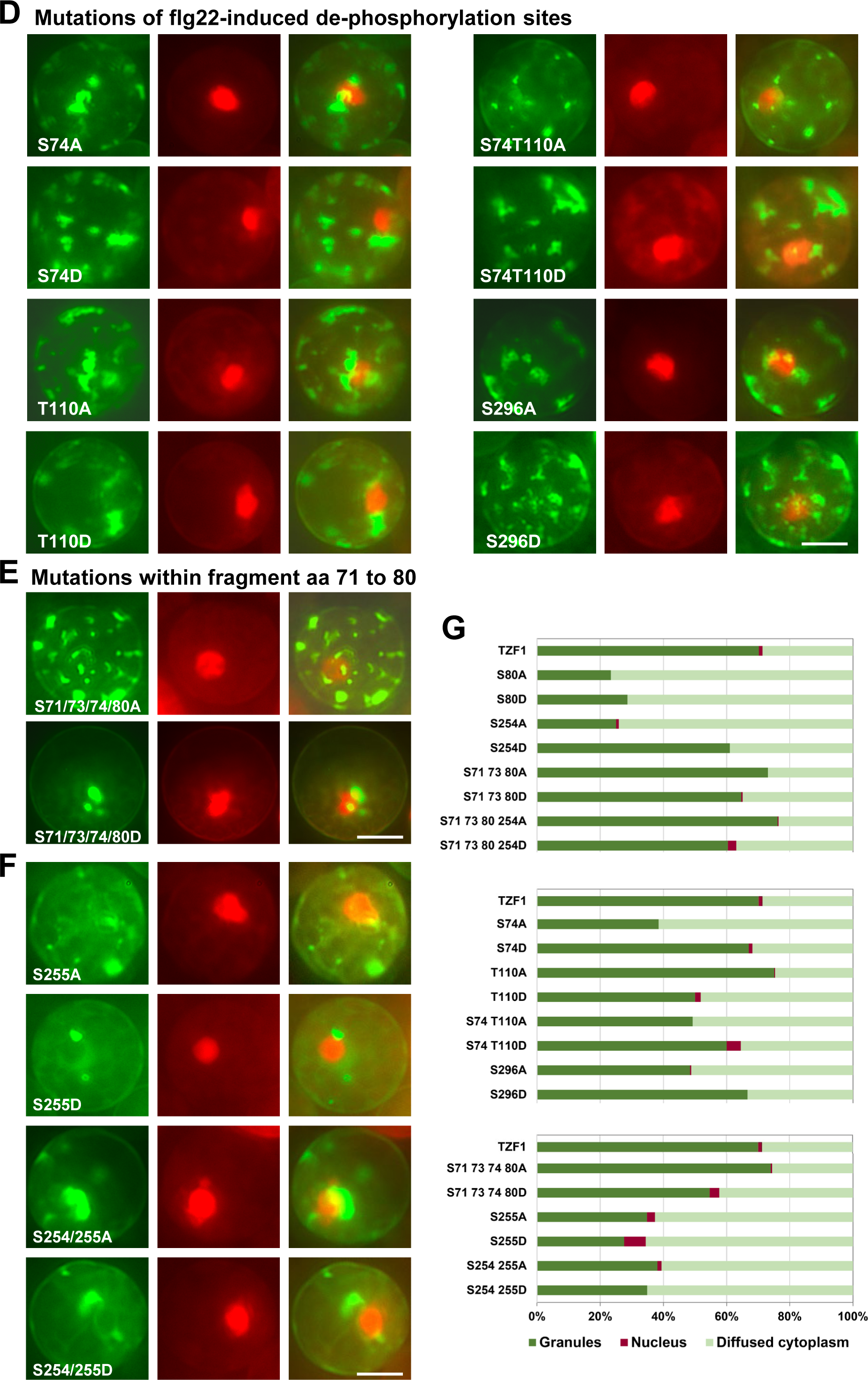
The effects of phosphorylation mutations on TZF1 cytoplasmic granule assembly. **(A)** TZF1-3FLAG-GFP fusion protein is mainly localized in cytoplasmic condensates/granules (#1) in *Arabidopsis* protoplasts. A small percentage of the cells displays diffused cytoplasmic (#2), nuclear (#3) or nucleus-like (#4) pattern. NLS-RFP, a marker for nuclear proteins. Scale bar= 10 μm. **(B)** TZF1 completely co-localized with SG marker UBP1b. **(C-F)** The subcellular localization patterns of TZF1-3FLAG-GFP with mutations on the residues phosphorylated (C), de-phosphorylated (D) upon flg22 treatment, a fragment with compact flg22-induced phosphorylated (S71, S73, and S80) or dephosphorylated (S74) residues (E), and a predicted MAPK phosphorylation residue (S255) and S254/255 double mutation (F). The S to A change represents phosphor-dead and S to D change represents phospho-mimicking mutation. Scale bar= 10 μm. **(G)** Quantitative analysis of TZF1-3FLAG-GFP subcellular localization patterns as shown in (A-F).

For flg22-induced phosphorylation sites (S71, S73, S80, and S254), most mutations caused reduction of TZF1 SGs, particularly S80, a conserved MPK phosphorylation site. Both S80A and S80D significantly reduced TZF1 SG assembly, in contrast to S254A with reduction and S254D with little change (Figures 6C and G). For flg22-induced de-phosphorylation sites (S74, T110, and S296), except for S74A showing slight decrease of SG assembly, none of the other mutants showed any significant change in the percentage of cells showing SG pattern (Figures 6D and G). Notably, the quadruple mutant of S71/73/74/80A caused an increase, whereas S71/73/74/80D caused a decrease in SG assembly (Figures 6D and G). Finally, the predicted MPK phosphorylation site mutation of either S255A or S255D along with the double mutants of S254/255A or S254/255D caused significant reduction in SG assembly (Figures 6F and G). These results indicated that protein phosphorylation plays a differential role in TZF1 SG assembly and there appears to be close interactions between phosphorylation events at different sites. For TZF1 protein accumulation, flg22-induced phosphorylation site phosphor-dead mutations (S80A, S254A, S71/73/80A, S71/73/80/254A) caused increase (Supplementary Figure 6A), whereas the corresponding phosphor-mimetic mutations (S to D change) caused slight decrease of TZF1 protein accumulation (Supplementary Figure 6B). For flg22-induced de-phosphorylation sites, the phosphor-dead mutations (S74A, S110A, S74/110A, S296A) did not appear to affect, while the corresponding phosphor-mimetic mutations (S to D change) caused significant reduction of TZF1 protein accumulation (Supplementary Figure 6). Finally, S255D and S254/255D mutants also caused drastic reduction of TZF1 protein accumulation. Together, these results suggest that de-phosphorylation causes increase, whereas phosphorylation causes decrease of TZF1 protein accumulation. For protein-protein interaction, neither flg22-induced phosphorylation nor de-phosphorylation mutations affect TZF1 interaction with MKK5 in SGs, albeit variations were observed in size and number of SGs (where TZF1 and MKK5 interacted) per cell (Supplementary Figure 7).

### The effects of 14-3-3 adaptor protein interaction sites

The IP-MS results indicated that TZF1 could potentially interact with two 14-3-3 adaptor proteins (Figure 1A). Using 14-3-3-Pred algorithm, 29 potential 14-3-3 protein interacting sites were predicted in TZF1. Among which S106, T168, T276, and S313 had the highest scores (Figure 7A). Because S106 fell within the RR domain and T168 fell within the TZF domain, site-specific mutations were made to test if these sites were important for TZF1 subcellular localization and protein-protein interaction. The S to A change (phosphor-dead to block 14-3-3 adaptor protein interaction) appeared to enhance the intensity of TZF1 SG signals (Figure 7B), although the percentage of cells with SGs remained largely unchanged (Figure 7C). Because S106 was identified as a phosphorylation site, the phosphor-mimetic mutant S106D was also examined. Interestingly, S106D significantly reduced the TZF1 SG assembly (Figures 7B-C), implicating that the interaction with 14-3-3 adaptor protein via S106 could potentially cause TZF1 SG disassembly via an as-yet-unknown mechanism. By contrast, T168A and S106T168A appeared to enhance TZF1 SG assembly (Figures 7B-C), suggesting that 14-3-3 interaction with TZF1 via T168 might play a negative role in TZF1 SG assembly. Immunoblot analysis revealed that the mutant proteins, including S106D, appeared to accumulate at higher levels than the WT TZF1 protein (Figure 7D). For protein-protein interaction, S106D, together with T168A and S106/T168A mutations appeared to enhance the interaction between TZF1 and MKK5 in SGs (Supplementary Figure 8). Therefore, it is likely that TZF1’s S106 plays a positive and T168 plays a negative role for 14-3-3 mediated TZF1-MKK5 interaction in SGs. The exact mechanism by which S106 and T168 affect TZF1-MKK5 interaction will be determined in future studies.

**Figure 7.**
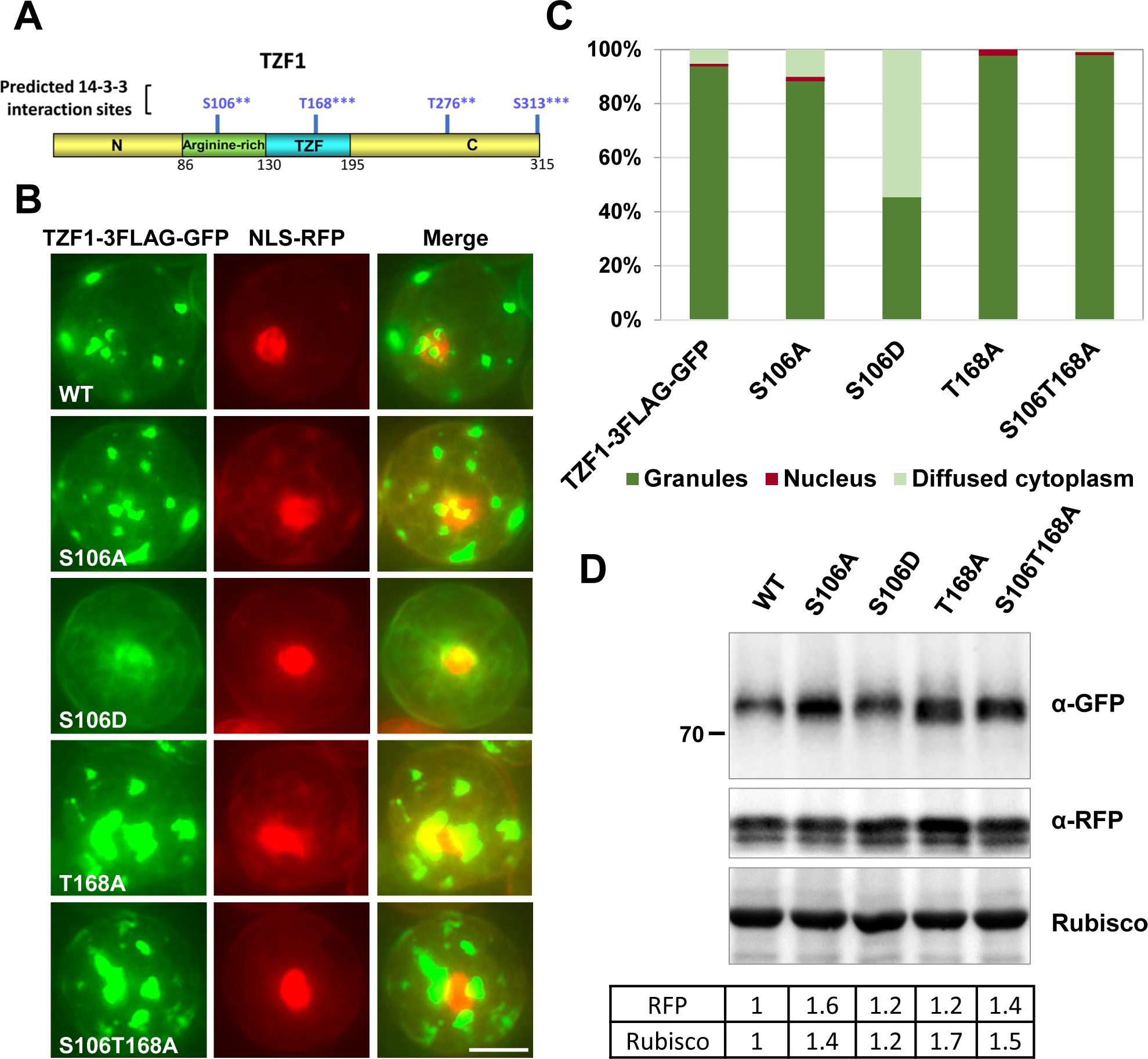
The effects of predicted 14-3-3 protein-protein interaction site mutations on TZF1 cytoplasmic granule assembly. **(A)** Four major 14-3-3 protein-protein interaction sites predicted by 14-3-3-Pred algorithm (https://www.compbio.dundee.ac.uk/1433pred). **(B)** Mutations (S/T to A) abolishing 14-3-3 interaction did not affect TZF1 localization to cytoplasmic granules, whereas S106D reduced cytoplasmic granule assembly. T168A appeared to enhance TZF1-GFP granule signal intensity. Scale bar= 10 μm. **(C)** Quantitative analysis of TZF1 subcellular localization patterns as shown in (B). **(D)** Immunoblot analysis of TZF1 (WT) and 14-3-3 interaction site mutations. Numbers in the table indicate normalized values of GFP/RFP and GFP/Rubisco, respectively.

### TZF1 protein accumulation is affected by KEG

Given KEG is associated with MKK4 and MKK5 (53) and all three components were also identified in our IP-MS analysis (Figure 2A), the functional relationship between TZF1 and KEG was investigated. Although KEG was reported to be localized in the trans-Golgi network and early endosomes (58), it could partially co-localize with TZF1 in cytoplasmic condensates (Figure 8A). TZF1 protein stability was then examined. In 7-day-old *CaMV35S:TZF1-GFP* (*TZF1-OX*) transgenic plants, TZF1 protein was extremely unstable—it almost completely disappeared after being treated by protein synthesis inhibitor cycloheximide (CHX) for just one hour. By contrast, its accumulation was restored by proteosome inhibitor MG115/132 cocktail. TZF1 accumulation could also be stabilized by PYR41, a ubiquitin-activating enzyme E1 inhibitor, or the combination of MG115/132 and PYR41 (Figure 8B). Consistently, TZF1 cytoplasmic granules were enhanced by MG132 in the root cells of *TZF1-OX* plants (Figure 8C). Immunoblot analysis indicated that TZF1-GFP accumulated at a higher level in the WT than in the *keg-4* gain-of-function mutant in either intact plant (Figure 8D) or in isolated protoplasts (Figure 8E). Furthermore, TZF1-GFP fluorescence signals were much stronger in the WT than in the *keg-4* in an Arabidopsis protoplast transient expression analysis (Figure 8F). Lastly, PYR41 inhibited SG assembly of TZF1(WT), but not the ubiquitination sites sextuple mutation (K to R change) TZF1^mU1-6^ (Figures 8G and H and 9A) due to inherited low number of SGs (described later). These results suggest that KEG might directly target TZF1 for ubiquitination-mediated degradation.

**Figure 8.**
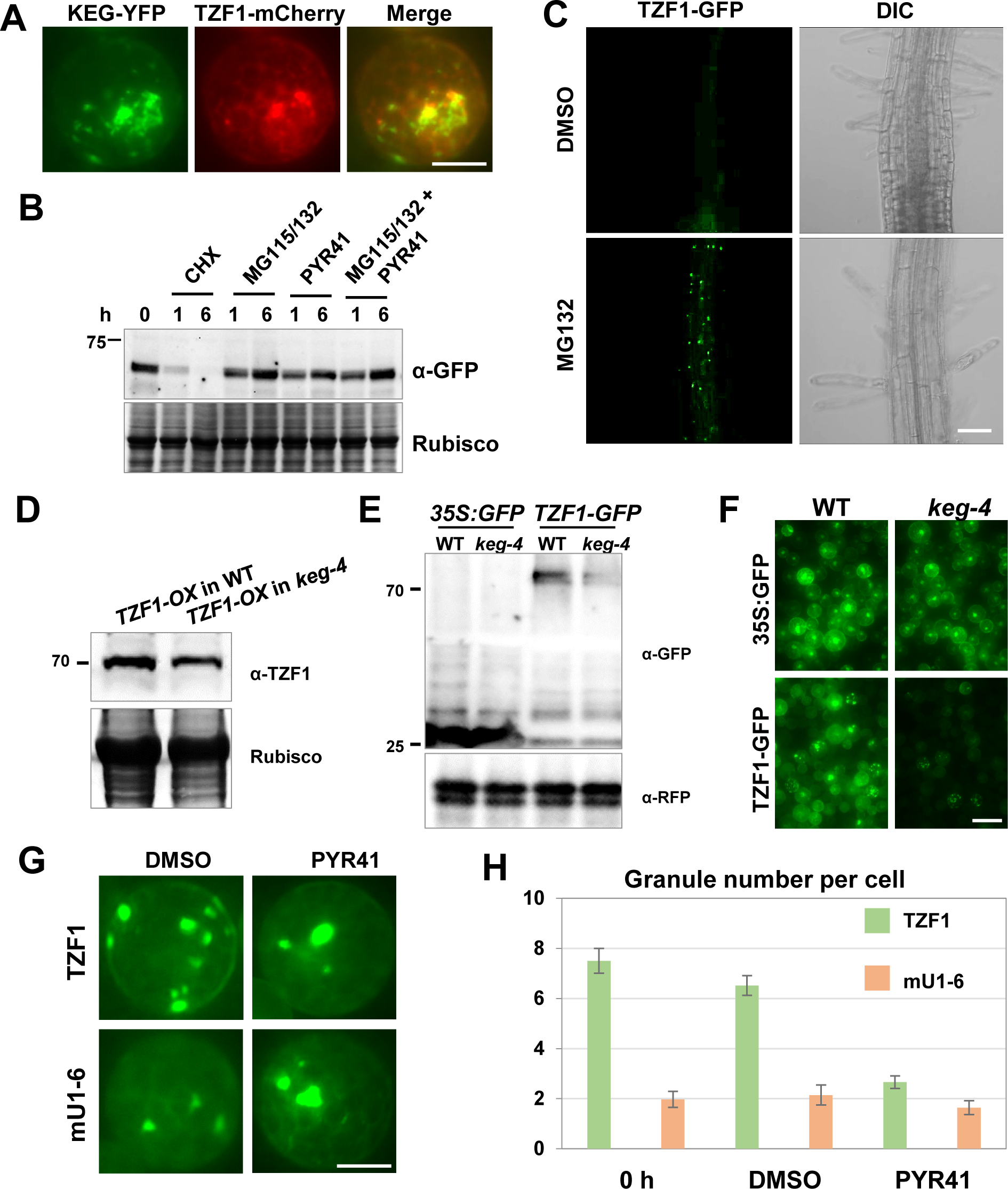
TZF1 accumulation is affected by KEG. **(A)** TZF1 is partially co-localized with KEG in cytoplasmic condensates. Scale bar= 10 μm. **(B)** TZF1 accumulation was blocked by protein synthesis inhibitor CHX and enhanced by proteosome inhibitor MG115/132 cocktail in 7-day-old *TZF1-OX* transgenic plants. **(C)** TZF1 cytoplasmic granules were enhanced by MG132 in the root cells of *TZF1-OX* transgenic plants. Scale bar= 20 mm. **(D)** Immunoblot analysis indicated that TZF1-GFP accumulated at a higher level in the WT than in the *keg-4* gain-of-function mutant. **(E)** Immunoblot analysis indicated that TZF1-GFP accumulated at a higher level in the WT than in the *keg-4* in *Arabidopsis* protoplasts. **(F)** TZF1-GFP signals were much higher in the WT than in the *keg-4* in *Arabidopsis* protoplasts. Scale bar= 30 mm. **(G and H)** TZF1 granule assembly was inhibited by PYR41, a ubiquitin-activating enzyme E1 inhibitor. The ubiquitination sites sextuple mutation (mU1-6) was relatively unchanged due to reduced number of TZF1 granules. Scale bar= 10 μm.

TZF1 was predicted to contain six putative ubiquitination sites by AraUbiSite tool (59) (Figure 9A). To investigate whether TZF1 is a substrate of KEG, we performed *in vivo* and *in vitro* ubiquitination assays to determine whether KEG could ubiquitinate TZF1. HA-tagged ubiquitin (HA-Ub) was co-expressed with TZF1-3FLAG-GFP in wild-type Arabidopsis leaf protoplasts. Ubiquitinated proteins were purified by IP with anti-HA antibody coated beads and then revealed by protein immunoblot analysis with anti-HA or anti-GFP antibody. Results showed that a potential ubiquitinated species was detected in TZF1, but not in the sextuple mutant TZF1^mU1-6^ (Figure 9B). Same IP experiment with the addition of PYR41 was also carried out. While the potential ubiquitinated TZF1 band was reproducible, the sample with the addition of PYR41 was too weak to determine if TZF1 ubiquitination could be abolished by PYR41 (Figure 9C). Because KEG is strongly self-ubiquitinated (60) (Figure 9D), ubiquitination blocked by PYR41 would stabilize KEG, hence enhance the degradation of TZF1. Next, we used recombinant GST-TZF1 and MBP-KEG (E3) to perform *in vitro* ubiquitination assays. High molecular-mass smear bands of TZF1 were detected in the presence of ubiquitin (Ub), E1 and E2, and MBP-KEG (E3 ligase) enzymes. However, the reactions without GST-TZF1 or MBP-KEG failed to produce any detectable upper smear bands of GST-TZF1. It was noted that KEG was strongly self-ubiquitinated (60) (Figure 9D). Together, these results indicate that TZF1 is likely ubiquitinated by KEG.

**Figure 9.**
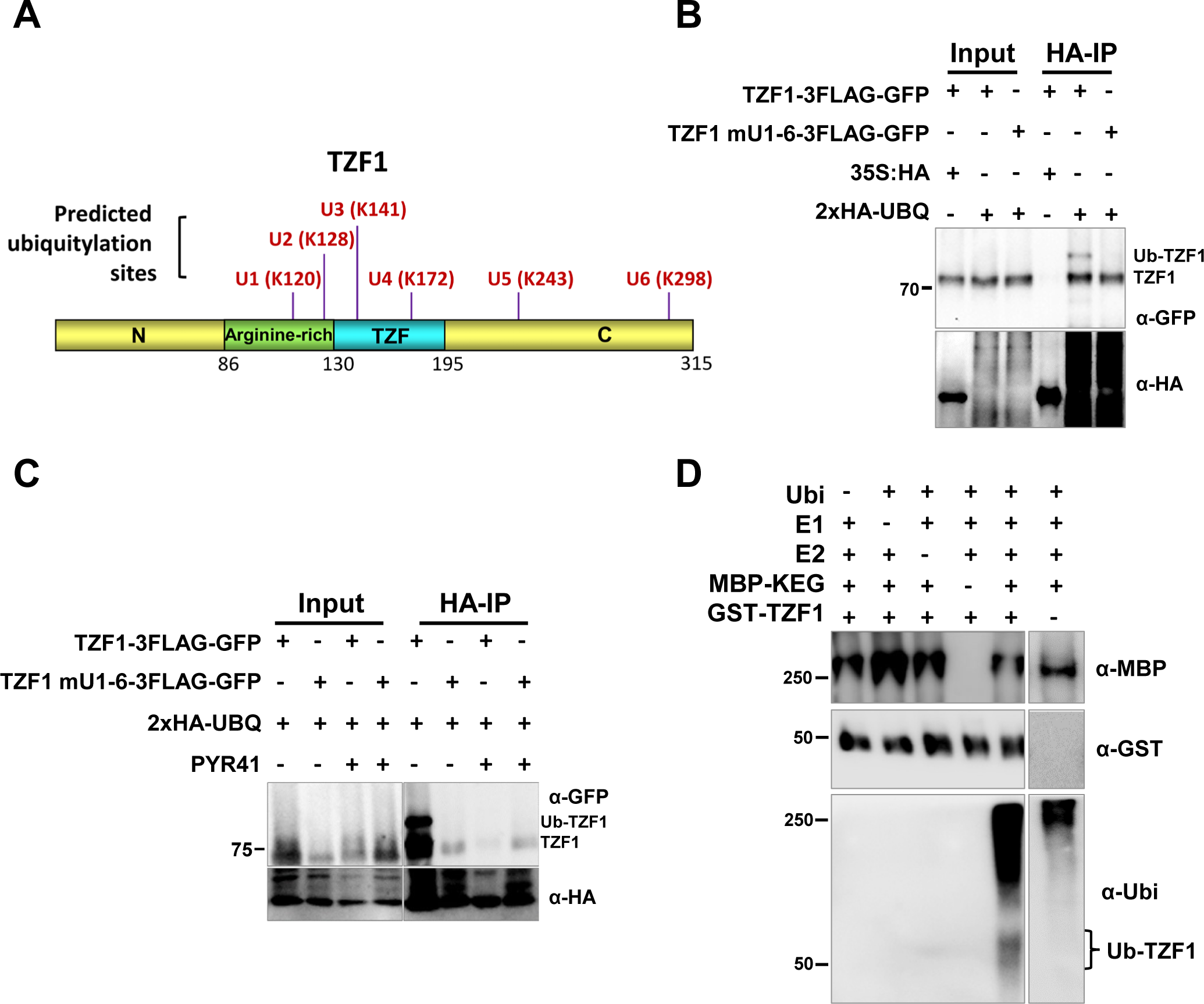
TZF1 is ubiquitinated *in vivo* and *in vitro*. **(A)** Schematic representation of ubiquitylated residues on TZF1 predicted by an online tool http://systbio.cau.edu.cn/araubisite. **(B)** TZF1 is ubiquitinated in vivo. Arabidopsis protoplasts were co-expressed with indicated constructs and IP was performed using anti-HA antibody and Western blot was carried out using anti-GFP antibody. (C) Same IP experiment with the addition of ubiquitin-activating enzyme E1 inhibitor PYR41 was carried out. **(D)** KEG ubiquitinates TZF1 *in vitro*. The in vitro reaction was carried out using recombinant E1, E2, and E3 (MBP-KEG) enzymes, ubiquitin, and GST-TZF1.

### Ubiquitination site mutations affect TZF1 SG assembly and protein-protein interaction with MKK5

To determine if ubiquitination of TZF1 affected SG assembly and protein-protein interaction with MKK5, site-directed mutagenesis was performed on predicted ubiquitination sites in TZF1 (Figure 9A). Intriguingly, except for TZF1^K172R^, all single and higher order mutations caused significant reduction of TZF1 SGs, particularly striking for TZF1^K120/128R^ (within RR motif) and TZF1^K141/172R^ (within TZF motif) double mutants. Note that both the percentage of cells with granule pattern as well as granule number per cell were reduced, except for TZF1^K172R^. Some large and single nucleus-like TZF1 SGs were observed in TZF1^K120/128R^, TZF1^K243R^, and TZF1^K120/128/141R^ mutants (Figures 10A-B), similar to the large SGs shown in (Figures 6A and B). For TZF1 interaction with MKK5, BiFC was conducted to access the subcellular localization of protein-protein interaction. Results showed that TZF1 mutants still interacted with MKK5 in SGs. Except for TZF1^K120/128R^, TZF1^K243R^, TZF1^K120/128/141R^, and TZF1^K120/128/141/172/243/298R^, most mutations caused reduced interaction based on the decreased number of SGs (where TZF1 and MKK5 interacted) (Supplementary Figure 9). For TZF1 protein accumulation, mutations within the RR-TZF domain (Figure 1B) appeared to reduce TZF1 accumulation. These included TZF1^K120/128R^, TZF1^K141R^, TZF1^K172R^, and TZF1^K141/172R^, except for TZF1^K120/128/141R^ with no effect on TZF1 accumulation. By contrast, mutations on C-terminal domain (TZF1^K243R^ and TZF1^K298R^) caused higher level of TZF1 accumulation, albeit not more than 30%. The higher order mutations (quadruple and up) caused significant decrease of TZF1 accumulation, due to unknown reasons (Figure 10C). Together, these results suggest that mutations of TZF1 ubiquitination sites have much more pronounced effects on SG assembly than protein stability control, perhaps due to the involvement of other unknown factors and interactions.

**Figure 10.**
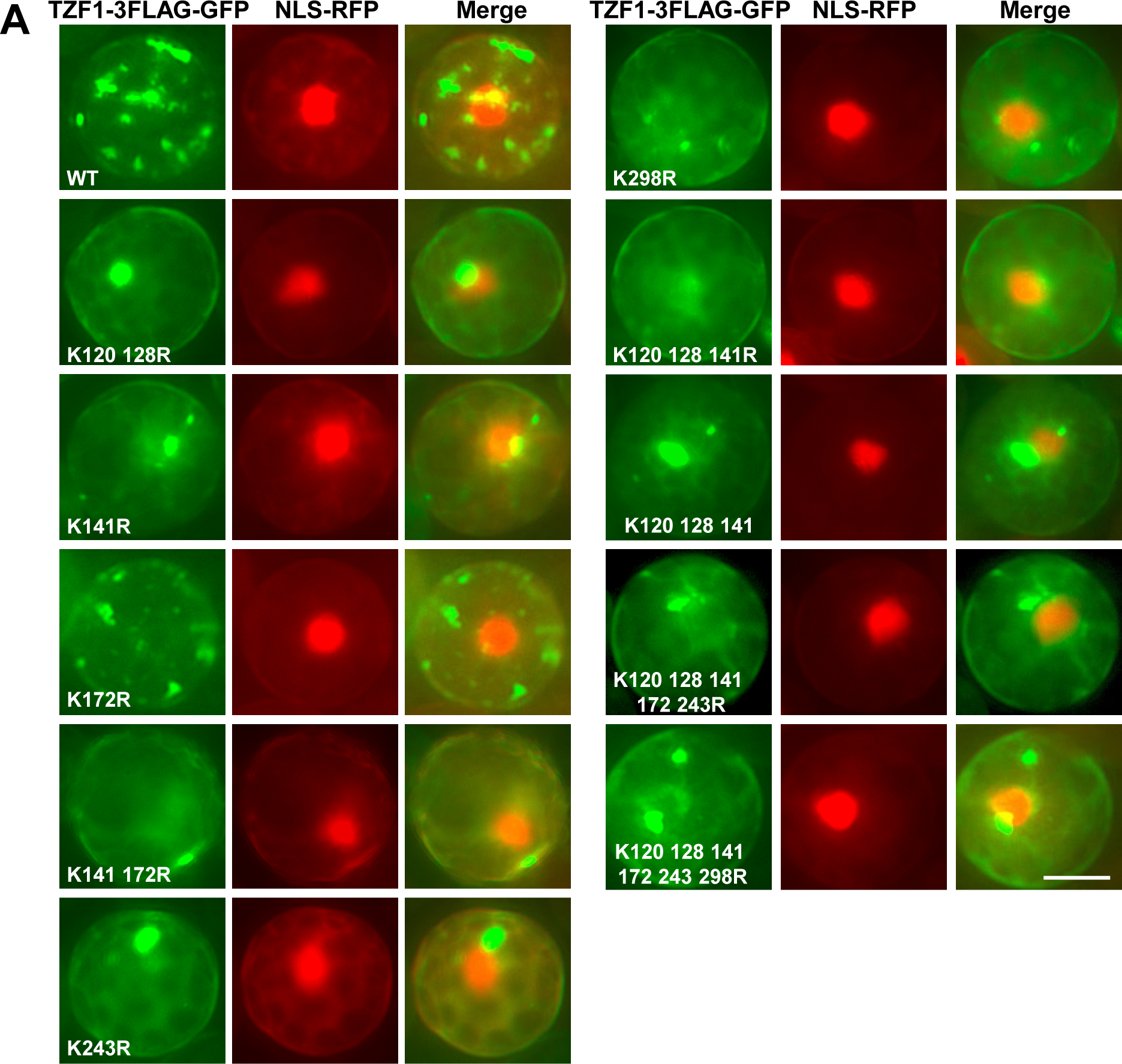
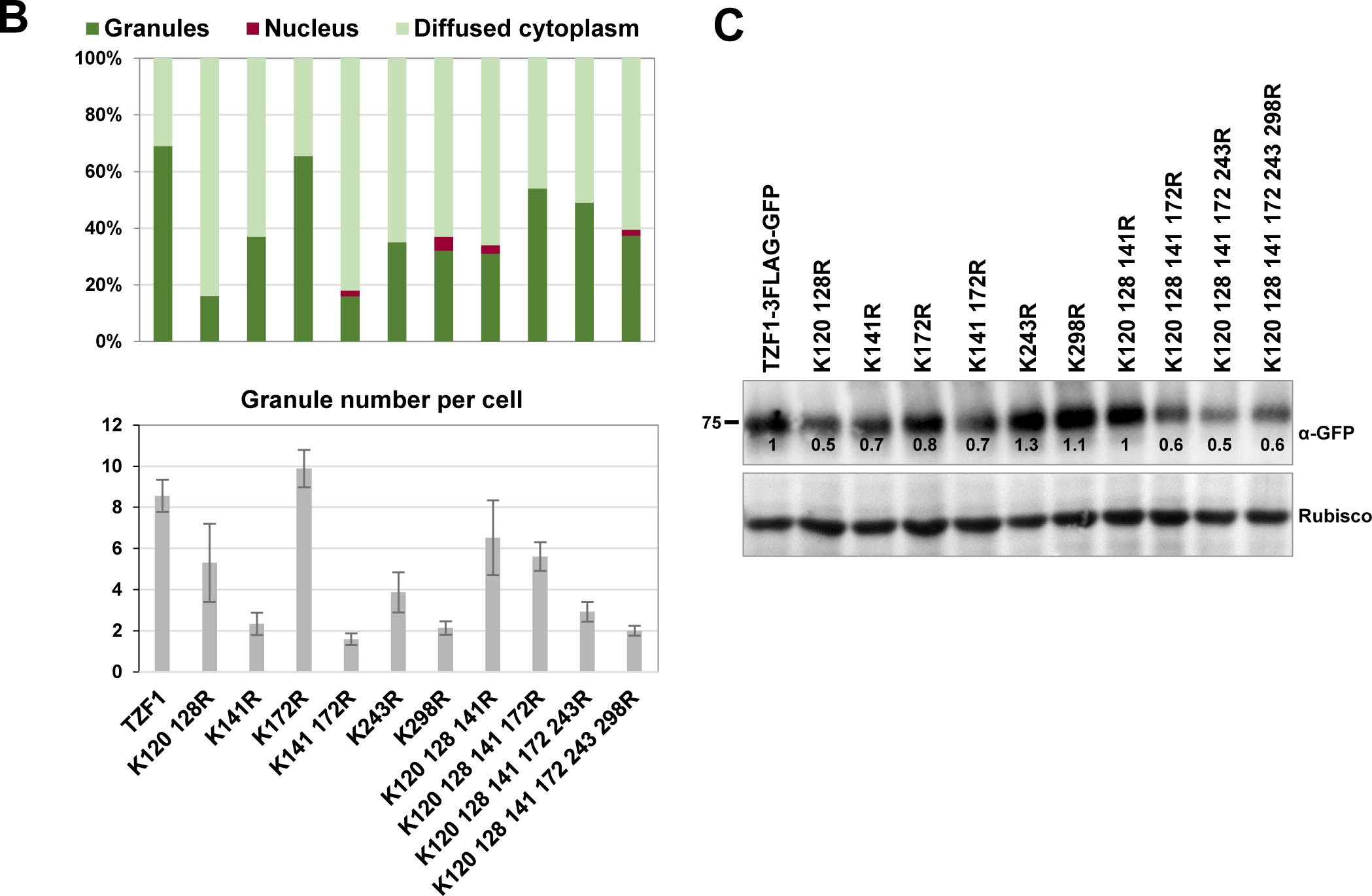
The predicted ubiquitylation site mutations abolished TZF1 cytoplasmic granule assembly in *Arabidopsis* protoplasts. **(A)** All mutations appeared to reduced TZF1 stress granule assembly. Scale bar= 10 μm. **(B)** Quantitative analysis of TZF1 subcellular localization patterns and granule number per cell as shown in (A). **(C)** Immunoblot analysis indicated that most ubiquitination site mutations reduced TZF1 protein accumulation.

## Discussion

Stress-induced RNP granules play pivotal roles in plant acclimation to various stresses and the class of SGs is conserved across different plant species (2). RNP granules regulate gene expression at the post-transcriptional and translational levels. The assembly/disassembly of RNP granules is intimately controlled by intra-and extra-cellular cues via signal transduction mainly mediated by PTMs of key protein components such as RBPs. The PTMs involved include but not limited to acetylation, arginine methylation, glycosylation, PARylation, phosphorylation, and ubiquitination. For example, the export mediators and translation regulators of stress-related mRNAs are recruited to SGs and phosphorylated by MAPKs, which results in changing the dynamics of SG-assembly and protein translation in animals (61).

### Phosphorylation and ubiquitination in TTP model

TTP is one of the most highly phosphorylated proteins in animals. To date, nearly 50 phosphorylation sites have been identified in mouse mTTP and human hTTP, respectively, in which it contains merely 319 and 326 amino acids (62). The stability and subcellular localization of TTP and its target mRNAs are tightly regulated by PTMs. The classical model of TTP-mRNA regulation is well established (Supplementary Figure 10A)—in the unstimulated condition, TTP is localized in the nucleus and SGs and both TTP and target mRNAs are labile. In the presence of proinflammatory stimulus, p38^MAPK^-MAPKAP kinase-2 (MK2) pathway is activated to phosphorylate TTP at S52 and S178 outside of TZF motif hence triggering 14-3-3 adaptor protein interaction. These events result in the exit of TTP from SGs to cytoplasm and stabilization of both TTP and target mRNAs (41,55).

With years of continuing research, layers of complexity have been added to this model (Supplementary Figure 10A). The N-terminal domain of TTP is phosphorylated by an unusual kinase MEKK1 (MAP triple kinase 1) and then the MEKK1 binding partner E3 ubiquitin ligase TRAF2 (TNF receptor-associated factor 2) deposits K63-linked ubiquitin chains onto five lysine residues within central TZF motif. The progressive decrease of TTP phosphorylation and increase of ubiquitination leads to the reduction of Nuclear Factor-kappaB (NF-kB) activity (pro-cell survival), whereas induction of the c-Jun N-terminal kinase (JNK) pathway (pro-cell death) (63). Curiously, the N-terminal domain of TTP also interacts with pyruvate kinase M2 (PKM2, typically a glycolysis enzyme) hence triggering p38^MAPK^-MK2 mediated phosphorylation, ubiquitination of TTP, reduction of target mRNA turnover, and impairment of cell viability in breast cancer cells (64).

In contrast to above mentioned non-degradative K63-mediated ubiquitination, TTP stability is also controlled by K48-mediated ubiquitination and degraded by 26S proteosome (Supplementary Figure 11A). HECT, UBA, and WWE domain-containing protein 1 (HUWE1) is a giant E3 ubiquitin ligase that regulates numerous substrates involved in signal transduction of cellular stress responses, cell growth, and apoptosis. A recent genetic analysis revealed that HUWE1 promotes the interaction between TTP C-terminal domain (aa 234-259) and protein phosphatases PP1 and PP2 and inhibits p38^MAPK^-MK2/JNK/ERK activities, therefore resulting in dephosphorylation of TTP (within aa 259-279) and leading to an activation of an unknow E3 ligase to deposit K48 ubiquitin chains onto the TZF motif to destabilize TTP protein. This pathway represents the late phase (3-16 h) of the pro-inflammatory stimulus-induced response (65).

### Phosphorylation

Similar to that in animals (1), plant SGs are formed via high local concentration and multivalent interaction of RNPs, where RBPs and signal transduction components such as kinases and phosphatases are enriched (2). Kinase signaling has an intimate relationship with SGs—either the assembly of SGs is dependent on kinase signaling or certain kinases themselves containing IDRs that could act as scaffolds to mediate SG assembly. SGs could serve as hubs to sequester kinases, cofactors, and substrates to spatiotemporally facilitate kinase signaling on client proteins of SGs (66). For example, protein kinase R is recruited to SGs by a core component G3BP1 to become active. By contrast, the yeast kinase Sky1 is recruited to SGs by its PrLD to phosphorylate other SG components thereby enhancing SG disassembly, thus acting as a negative feedback mechanism for SG growth (66). In plants, little is known about the roles of protein phosphorylation on the assembly of and protein-protein interaction with in SGs. Although numerous reports have demonstrated the central roles of scaffold proteins and PTMs in nucleating and promoting SG assembly, none except G3PB-deficient mutants completely failed to form SGs in non-plant systems, and surprisingly this has not been observed in plant system (2).

In this report, we demonstrate the interaction and localization of MAPK signaling components in TZF1-associated SGs. These results are strongly supported by a previous protein interactome analysis indicating that MPK3, MKK4, and MKK5 were found in various SG proteomes (2). We also unequivocally demonstrated that TZF1 is an IDR-containing SG component likely playing a key role in RNA metabolism and signal transduction (67). Our study made a step further to show that MPKs and MKKs are recruited by TZF1 to SGs and TZF1 is phosphorylated by MPK3/6 (Figures 2-4 and Supplementary Figures 2 and 4). The TZF1-MPK/MKK interaction in SGs is further substantiated by our IP-MS results in which conserved SG markers RH6/8/12 are also found in TZF1 protein complex (Figure 2A). We then fine mapped the phosphorylation sites of the TZF1 using LC-MS/MS to reveal 10 potential residues, among which S80 and S254 are associated with MPK phosphorylation signature motif SP, and S106 and T276 are within predicted 14-3-3 adaptor protein binding sites (Figure 5).

Our comprehensive mutant studies have revealed that phosphorylation has differential effects on TZF1 SG assembly and protein-protein interaction with MKK5 in SGs, depending on the location and phosphorylation status of the amino acids. As mentioned earlier, it was not surprising that none of the single or higher order mutations of either S to A or S to D changes could abolish SG assembly, given no prior examples could be found in plant systems. However, a significant reduction of SG assembly was found in the mutations of S80 and S254/255 (Figure 6), the two predicted MAPK phosphorylation sites. It is currently unclear though why both S to A and S to D changes resulted in similar SG reduction, given there has been no prior research on TZF1 phosphorylation, let alone the roles of the two specific amino acid residues in SG assembly. We speculated that the phosphorylation status of these residues is tied to a feedback regulatory loop of SG homeostasis. Disruption of the balance of such loop controlled by reversible phosphorylation could lead to the disassembly of SGs (Figure 13).

Another striking result we have obtained is the relationship between phosphorylation and TZF1 protein accumulation. Although standard protein half-life analyses were not performed, the S to D mutations of almost all residues tested result in lower protein accumulation (Supplementary Figure 6). This is in sharp contrast with the animal TTP model (Supplementary Figure 10A) in which phosphorylated TTP is more stable (41,55). Moving forward, it is imperative to unbiased determine and unravel the mechanism underpinning phosphorylation mediated control of TZF1 protein and target mRNAs stability.

### 14-3-3 adaptor protein interaction

The recruitment of 14-3-3 adaptor protein by MAPK signaling could have a significant impact on TZF1 function because 14-3-3 proteins could mask the IDRs of TZF1 and reduce its multivalency hence acting as inhibitors of TZF1-mediated SG assembly (66) (Supplementary Figure 10). In this report, we confirm the role of IDR in SG assembly because deletion of either or both IDRs in TZF1 almost eliminated SG assembly completely (Figures 1B-C). In addition, the deletion of IDRs from TZF1 also changed the pattern of protein-protein interaction with MKK5 from random granular to few large coalesced SGs attached to the nucleus (Results not shown and Figure 13). This could be due to the compositional change in protein, RNA, or both in SGs. Furthermore, phosphorylation of a potential 14-3-3 interaction site (S106D) significantly reduced TZF1 SG assembly (Figure 7). Reduced SG assembly could be interpreted by the masking of TZF1’s IDRs by 14-3-3 adaptor protein via phosphorylation dependent protein-protein interaction and thus reducing TZF1’s ability to recruit other components for SG assembly.

### Ubiquitination

Ubiquitination is generally considered as a switch that destines protein for degradation by 26S proteosome. However, protein mono-ubiquitination and K63 type poly-ubiquitination are non-proteolytic signals that serve as means in controlling other cellular processes such as protein-protein interaction and protein phosphorylation, as has been intensively investigated in the NF-kB pathways. In animals, ubiquitination also regulates the activation of MAP kinases in immune and inflammatory pathways (68). In another scenario, the K63-ubiquitination in the cells is required for DCP1a phosphorylation, decapping and mRNA degradation of prototypical inflammatory genes, and most remarkably the assembly of decapping factors into P-bodies. Curiously, mutation of all six ubiquitin acceptor lysine residues (K520-577R) at the C-terminal of DCP1a increased the number but reduced the size of DCP1a-associated P-bodies, illustrating a multifaceted regulation of ubiquitination on the dynamics of P-body assembly (69).

Conversely, kinases could act as sensors for the PTM events taking place in LLPS-mediated condensates. For example, some kinases (e.g., TANK-binding kinase 1—TBK1) can sense ubiquitin and be recruited and activated (e.g., through oligomerization due to elevated local concentration) in the condensates enriched with ubiquitin-tagged mis-folded proteins. This feed forward pathway can promote condensate growth and recruit additional polyubiquitin-tagged proteins to eventually trigger the participation of aggrephagy machinery (via TBK1 phosphorylation of aggrephagy receptors) to clear toxic protein aggregates that cause degenerative diseases such as amyotrophic lateral sclerosis (ALS) (66). Curiously, it was reported that *Drosophila* TTP homolog dTIS11 protein was unstable and it could be degraded by 20S proteosome in a phosphorylation dependent but ubiquitination and ATP independent manner. This unique mechanism was attributed to the IDRs at both N-and C-terminus of dTIS11 that render the protein susceptible to degradation by 20S proteosome (70).

In plants, an activator of salicylic acid induced systemic acquired resistance NPR1 (nonexpresser of PR genes 1) is recruited to the cellular condensates to trigger a partner E3 ubiquitin ligase mediated ubiquitination of other proteins in the condensates to enhance cell survival (71). In this report, we show that TZF1 is ubiquitinated by E3 ubiquitin ligase KEG (Figure 9). Intriguingly, mutations of predicted ubiquitination sites of TZF1 significantly reduce SG assembly and some mutations trigger the formation of large coalesced SGs directly in contact with the nucleus (Figures 10 and 13). It is well documented that RNP granules can undergo homotypic or heterotypic interaction to facilitate the assembly or larger granules. In general, SGs prefer to interact with themselves and two or more SGs can dock and form a larger condensate. By contrast, it is relatively rare for heterotypic docking of PBs with SGs to allow the exchange of RNPs including mRNAs (1). We propose that TZF1 ubiquitination facilitates homotypic interaction via docking and merging to form larger SGs. However, we cannot rule out the possibility of heterotypic interaction, given TZF1 could partially localize to PBs as well (48). It is currently unclear whether the composition or property of TZF1 granules would be changed during the formation of a single or multiple nucleus-like large condensate within the cells (Figure 6A). As was reported recently, RNAs were primarily degraded in smaller liquid-like PBs, whereas RNAs were mostly stable under heat shock condition when PBs increased in size and became more solid-like (72). In our study, we do not know the property, composition, and fate of the RNAs associated with TZF1 condensates, but we do observe in numerous occasions on the dynamic changes of the size and number of TZF1 condensates among various mutations (Figure 13), implicating that post-translational modification such as phosphorylation and ubiquitination might control TZF1 condensates’ ability in modulating mRNA metabolism. On the other hand, it is currently unclear how the large TZF1 SGs are connected to the nucleus. Perinuclear RNA granules such as germ granules (known as P-granules in *Caenorhabditis elegans*) are well characterized to be associated with nuclear pore complex (73). However, it could be up to a dozen of even-sized P-granules surrounding a single nucleus, which is quite different from what observed here the one to one ratio of a large TZF1 SG to the nucleus. The cause and biological significance of large coalesced SGs docked to the nucleus are important questions to be addressed in the future.

Another striking result in our study is the reduction of TZF1 protein accumulation resulted from mutations of ubiquitination sites (Figure 10C). We do not know if TZF1 is ubiquitinated via K48 or K63 ubiquitin chain. Given the results of protein immunoblot analysis we have obtained, it might be more likely that TZF1 is predominantly K63-ubiquitinated. Mutagenesis of ubiquitination sites is frequently used to validate ubiquitination targets. However, the results from using this approach can be difficult to interpret due to unintended changes of protein folding, protein-protein interaction, ATP/ubiquitin binding, and protein activities (68). Therefore, reduced TZF1 accumulation caused by ubiquitination site mutations could be a consequence of multiple reasons mentioned above. A deeper dissection on the mechanisms of TZF1 ubiquitination is required to address these important open questions in the future.

In summary, we have found that TZF1 recruits MAPK signaling components and an E3 ubiquitin ligase KEG to SGs (Figure 11). TZF1 is then phosphorylated by MPKs and ubiquitinated by KEG. In this process, we have found that Arabidopsis TZF1 is not less complicated than animal TTP, in terms of domain/structure and function (Supplementary Figure 10A). Phosphorylation, ubiquitination, 14-3-3 interaction, IDRs, and numerous regulatory elements throughout TZF1 might all have differential effects on RNP granule assembly, and protein-protein interaction with a key MAPK signaling component MKK5 in SGs (Figure 13). Given decades of intensive studies on mammalian TTP, our understanding on Arabidopsis TZF1 thus far appears to be in its infancy and one-dimensional (Supplemental Figure 10B). However, we believe our groundbreaking study has served as a gateway for the in-depth investigation in the future. Transgenic plants for all the constructs discussed here have been generated. Work is in progress to obtain homozygous lines. Moving forward, in-depth characterization of these transgenic plants is expected to gain more insights into plant stress granules dynamics in response to various cues. Because TZF family proteins are evolutionarily conserved not only in sequence and structure but also in expression pattern and function, much more intensive and intelligent studies are required to translate basic information into useful new tools for crop improvement.

**Figure 11.**
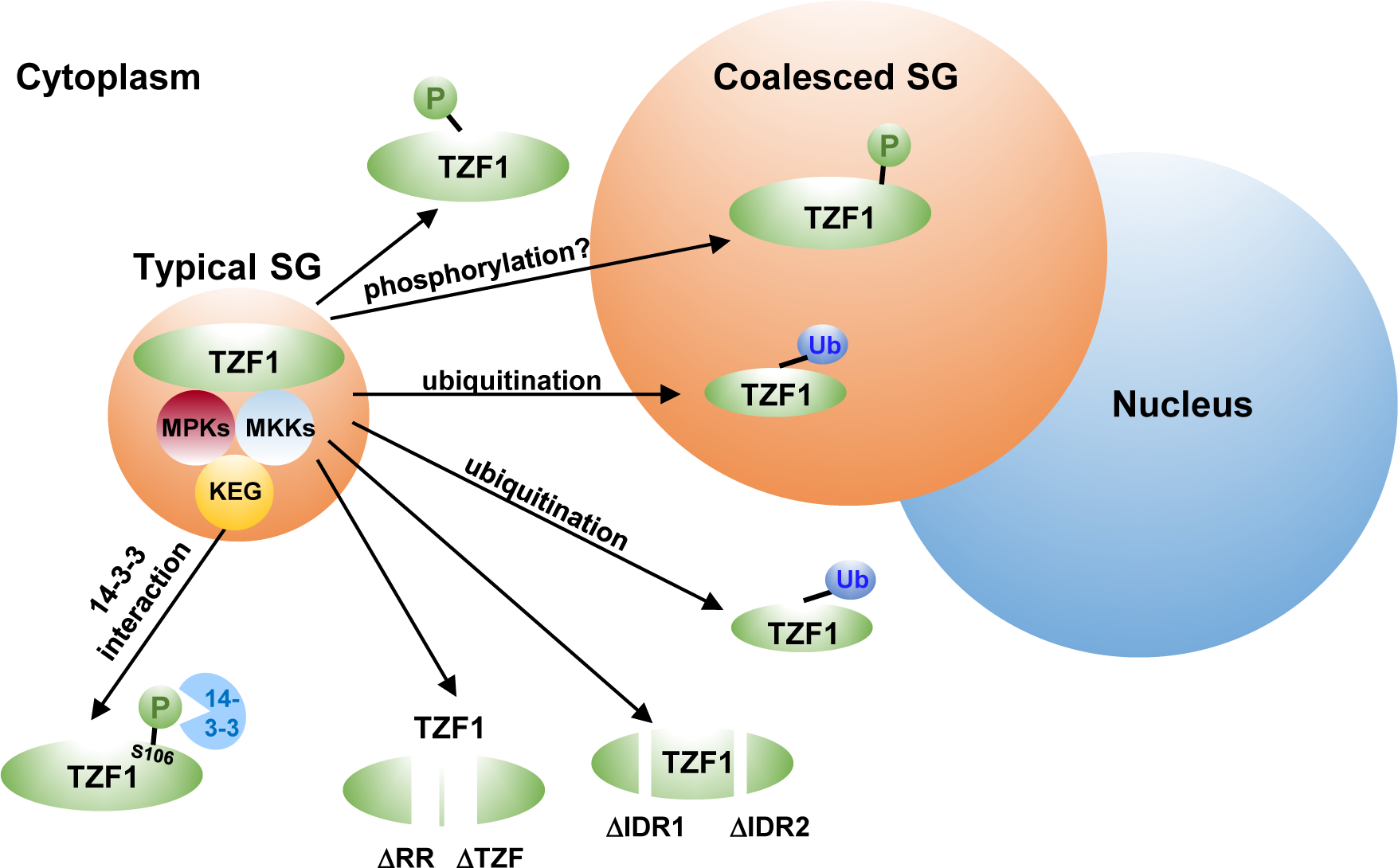
Effects of post-translational modifications on TZF1 SG assembly. *Arabidopsis* TZF1 recruits MAPK signaling components MPK3, MPK6, MKK4, and MKK5 and an E3 ubiquitin ligase KEG to SGs. TZF1 is phosphorylated by MPK3/6 and ubiquitinated by KEG. Depending on the position and status of the phosphorylation and ubiquitination modifications, TZF1 subcellular localization can be changed from a typical SG pattern to SG disassembly to become cytoplasmic pattern or to the formation of one or more coalesced large SGs attaching to the nucleus. Deletion of IDR, RR, TZF motif and phosphorylation-induced 14-3-3 interaction at TZF1 (S106) can also result in the reduction of SG assembly.

## Materials and Methods

### Plant materials and growth conditions

*Arabidopsis thaliana* ecotype Columbia (Col-0) was used in this study. The *keg-4* mutant (CS67951) was obtained from the Arabidopsis Biological Resource Center (ABRC). WT, *keg-4*, and transgenic plants were grown in a growth chamber at 22℃ with a photoperiod of 16-h light/8-h dark.

### Molecular cloning and generation of transgenic plants

The coding sequence (CDS) of TZF1, MKK4, and MKK5 were cloned into the pENTR™/D-TOPO^®^ vector. All constructs were subcloned into the Gateway^®^ destination binary vector with C-terminal GFP tag by using the LR recombination reaction and then transformed into WT plants by the floral dip method. The constructs used for phosphorylation and ubiquitination mutant analysis were cloned into a modified pBlueKS+ plasmid with LR recombination sites as a Gateway destination vector (30).

### Yeast two-hybrid (Y2H) assay

The CDS of TZF1 was cloned into the pGBKT7 vector and the CDS of MKK4, MKK5, MPK3, and MPK6 were cloned into the pGADT7 vector. Pairs of pGBKT7 and pGADT7 plasmid were co-transformed into the yeast strain AH109 following the Matchmaker^TM^ GAL4 Two-Hybrid System instructions (Clontech). Primary transformants were selected on synthetic drop-out (SD) medium lacking Trp and Leu and confirmed again by colony PCR before growing on SD medium lacking Ade, His, Trp, and Leu.

### Protoplast transient expression and BiFC assays

For transient expression assay in *Arabidopsis* protoplasts, TZF1, MKK4, MKK5, MPK3, and MPK6 CDS were cloned into the pENTR™/D-TOPO^®^ vector and then subcloned into the Gateway^®^ destination vector with C-terminal GFP tag by using the LR recombination reaction. For BiFC, the CDS of TZF1 was cloned into pA7-YN (containing N-terminal half of YFP) vector and the CDS of MKK4, MKK5, MPK3, and MPK6 were cloned into pA7-YC (containing C-terminal half of YFP) vector (74). Plasmid pairs were co-transformed into *Arabidopsis* protoplasts.

### Co-IP assay

Total proteins were extracted from *Arabidopsis* protoplasts co-expressing *TZF1-2xHA* with *GFP-MPK3*, *GFP-MPK6*, *MKK4-GFP*, *MKK5-GFP* or free *GFP*. Extracted proteins were then incubated with equilibrated GFP-trap beads (Chromotek) at 4°C for 2 h under gentle agitation, followed by 3 times of washing with wash buffer (100 mM Tris pH 8.0, 150 mM NaCl, 5 mM EDTA, 10 mM DTT, 0.1% NP-40). Immunoblots were performed using α-GFP (Roche) or α-HA antibodies (Roche).

### *In vivo* ubiquitination assay

*Arabidopsis* protoplast samples were co-transformed with *2xHA-UBQ* and the GFP-tagged genes of interest and incubated overnight at room temperature followed by a 2-h treatment with 50 μM MG132. After homogenization in 100 μl of IP buffer (100 mM Tris pH 8.0, 150 mM NaCl, 5 mM EDTA, 10 mM DTT, 0.1% NP-40, protease inhibitor cocktail), the GFP-tagged proteins were immunoprecipitated by incubating the extracts with 15 μl of anti-HA magnetic beads (Thermo Scientific) for 2 h at 4°C with gentle shaking. The anti-HA magnetic beads were collected and washed 3 times with wash buffer (100 mM Tris pH 8.0, 150 mM NaCl, 5 mM EDTA, 10 mM DTT, 0.1% NP-40). Immunoblots were performed using α-GFP (Roche) or α-HA antibodies (Roche).

### *In vitro* ubiquitination assay

The *in vitro* ubiquitination reaction was performed in a 30 μl mixture containing 200 ng E1 enzyme (BB-E-304-050, Boston Biochem), 200 ng E2 enzyme (BB-E2-616-100, Boston Biochem), 5 mg His-ubiquitin (BB-U-530, Boston Biochem), 2 mg purified MBP-KEG fusion protein (as E3 enzyme), and GST-TZF1 fusion protein in a reaction buffer that contains 50 mM Tris–HCl [pH 7.6], 2 mM DTT, 5 mM MgCl_2_, and 2 mM ATP. After 1 h incubation at 30°C in Eppendorf Thermomixer, the reactions were stopped by adding SDS-PAGE sample buffer. Ubiquitinated proteins were detected using ubiquitinh antibody. MBP-KEG was detected by anti-MBP monoclonal antibody and GST-TZF1 was detected by anti-GST monoclonal antibody.

### Identification of TZF1 phosphorylation sites by mass spectrometry

To identify TZF1 phosphorylation sites, TZF1-HA was expressed in *Arabidopsis* protoplasts (concentration of 2 × 10^5^/ml) for 12 h and treated with or without 0.1 μM flg22 for 15 min. Ten mL protoplasts were used to immunoprecipitate TZF1-HA proteins from mock and flg22-treated samples, respectively. Protoplasts were then lysed with lysis buffer (20 mM Tris-HCl, pH 7.5, 100 mM NaCl, 10% glycerol, 0.5 Triton X-100, 1 mM EDTA, 2 mM DTT, 2 mM NaF, and 2 mM Na_3_VO_4_, and 1x protease inhibitor EDTA-free cocktail) and immunoprecipitated with α-HA magnetic beads (Thermo Fisher). The immunoprecipitated products were separated by 10% SDS-PAGE and stained with GelCode Blue Stain Reagent (Thermo Fisher) for 2 h at 23°C. The TZF1-HA bands were sliced, trypsin-digested, and phospho-peptides were subjected to LC-MS/MS analysis using an Orbitrap QE LC-MS/MS system (Thermo Scientific) at the proteomics core facility of UT Southwestern Medical Center. The MS/MS spectra were analyzed with Mascot software, and the identified phosphor-peptides were manually inspected to ensure the accuracy of phosphorylation sites detection.

## Author Contributions

S.-L.H. and J.-C.J. conceived and designed the experiments; S.-L.H. performed most of the experiments, X.W. performed *in vitro* ubiquitylation assay, L.K. identified TZF1 phosphorylation sites by mass spectrometry; S.-L.H. and J.-C.J. wrote the manuscript with input from all authors. L.W., S.K., P.H., L.S., and Y.W. provided suggestions and comments for the project.

## Acknowledgements

We thank Dr. Roger W. Innes (Department of Biology, Indiana University, Bloomington, Indiana) for providing the KEG-YFP plasmid.

## Supplementary data

**Supplementary Figure 1. The effects of intrinsically disordered regions (IDRs) and RR-TZF motif on TZF1 protein accumulation.**

Arabidopsis protoplast transient expression system was used to generate indicated samples for Western blot analysis. *CaMV35S:NLS-RFP* was co-expressed as an internal control. Rubisco band was used as loading control.

**Supplementary Figure 2. TZF1 interacts with MKKs and MPKs by BiFC.**

TZF1 interacts with MKK4 **(A)**, MKK5 **(B)**, MPK3 **(C)**, and MPK6 **(D)** in BiFC analyses. YN/YC: N-and C-terminal fragment of yellow fluorescence protein, respectively. Shown are confocal microscopy images revealing that the protein-protein interactions take place in cytoplasmic condensates that partially co-localize with PB marker DCP1-mCherry, but completely co-localize with SG marker UBP1b-mCherry. NLS-RFP, a marker for nuclear proteins. Scale bar= 10 μm.

**Supplementary Figure 3. Controls for bimolecular fluorescence complementation (BiFC) analysis.**

**(A)** TZF1 BiFC constructs were reciprocally co-expressed with corresponding empty vectors. Various C-terminal YFP fusion constructs were co-expressed with the empty vector containing N-terminal YFP in Arabidopsis protoplasts. **(B)** MKK5 fused with C-terminal YFP was co-expressed with various negative control proteins fused with N-terminal YFP in BiFC analysis using Arabidopsis protoplast transient expression system. Scale bar= 10 μm.

**Supplementary Figure 4. Co-localization of MAP kinase signaling components with DCP1 and UBP1b.**

The MAP kinase signaling components MKK4 **(A)**, MKK5 **(B)**, MPK3 **(C)**, and MPK6 **(D)** are not co-localized with PB marker DCP1-mCherry, but completely co-localized with SG marker UBP1b-mCherry. Scale bar= 10 μm.

**Supplementary Figure 5. Three-dimentional rotating view of a large coalesced TZF1 SG (in green fluorescence) attached to the nucleus (in red).** Scale bar= 5 μm.

**Supplementary Figure 6. Results of immunoblot analysis showing the accumulation of TZF1 phosphorylation sites mutants as shown in Figure 5**.

Supplementary Figure 7. MKK5 interacts with TZF1 WT and mutants with flg22-induced phosphorylation site mutations (A) and with flg22-induced de-phosphorylation site **(B)** in cytoplasmic condensates in BiFC analyses. Scale bar= 10 μm.

**Supplementary Figure 8. The effects of TZF1 predicted 14-3-3 protein-protein interaction sites on TZF1 interaction with MKK5 in BiFC analysis.**

TZF1 WT and mutants of 14-3-3 protein-protein interaction sites interact with MKK5 in cytoplasmic condensates in BiFC analyses. The S106A mutation appeared to cause reduction of BiFC signal. Scale bar= 10 μm.

**Supplementary Figure 9. The effects of TZF1 predicted ubiquitination sites on protein-protein interaction with MKK5.** TZF1 WT and mutants of ubiquitination site mutations interact with MKK5 in cytoplasmic condensates in BiFC analyses. The K120/128R and K243R mutations appeared to cause reduction of BiFC signals. Scale bar= 10 μm.

**Supplementary Figure 10. Comparison of phosphorylation-and ubiquitination-mediated regulatory pathways between mouse TTP and *Arabidopsis* TZF1 models.**

**(A)** Pro-inflammatory stimulus activates p38^MAPK^-MK2 pathway to phosphorylate TTP at S52 and S178 hence triggering 14-3-3 adaptor protein interaction. These events result in the exit of TTP from SGs to cytoplasm and stabilization of both TTP and target mRNAs (41,55). The N-terminal domain of TTP is phosphorylated by MEKK1 and then the E3 ubiquitin ligase TRAF2 deposits K63-linked ubiquitin chains onto TZF motif. The progressive decrease of TTP phosphorylation and increase of ubiquitination leads to the reduction of NF-kB activity (pro-cell survival) but induction of the JNK pathway (pro-cell death) (63). The N-terminal domain of TTP also interacts with PKM2 hence triggering p38^MAPK^-MK2 mediated phosphorylation, ubiquitination of TTP, and reduction of target mRNA turnover ability (64). In the late phase (3-16 h) of the pro-inflammatory stimulus-induction, HECT, UBA, and HUWE1 promotes the interaction between TTP C-terminal domain (aa 234-259) and protein phosphatase PP1 and PP2 and inhibits p38^MAPK^-MK2/JNK/ERK activities, therefore resulting in dephosphorylation of TTP (aa 259-279) and leading to an activation of an unknow E3 ligase to deposit K48 ubiquitin chains onto the TZF motif to destabilize TTP protein (65). **(B)** MAMPs such as flg22 activates MAPK signaling cascade to induce plant defense response. TZF1 recruits MKK4/5, MPK3/6, and an E3 ubiquitin ligase KEG to SGs where TZF1 is phosphorylated by MPK3/6. Via self-phosphorylation or phosphorylation by MPKs (?) (60), KEG is self-ubiquitinated and it also ubiquitinates MKK4/5 (53) and TZF1. The phosphorylation of S106 and T168 (?) can potentially trigger 14-3-3 adaptor protein interaction and affect TZF1 protein stability, protein-protein interaction, and SG assembly.

